# Zebrafish Ski7 tunes RNA levels during the oocyte-to-embryo transition

**DOI:** 10.1101/2020.03.19.998716

**Authors:** Luis Enrique Cabrera Quio, Alexander Schleiffer, Karl Mechtler, Andrea Pauli

## Abstract

Post-transcriptional mechanisms are crucial for the regulation of gene expression. These mechanisms are particularly important during rapid developmental transitions such as the oocyte-to-embryo transition, which is characterized by dramatic changes to the developmental program in the absence of nuclear transcription. Under these conditions, changes to the RNA content are solely dependent on RNA degradation. Although several mechanisms that promote RNA decay during embryogenesis have been identified, it remains unclear which cellular machineries contribute to remodeling the maternal transcriptome during the oocyte-to-embryo transition. Here, we focused on the auxiliary 3’-to-5’ degradation factor Ski7 in zebrafish as its mRNA peaks during this time frame. Homozygous *ski7* mutant fish were viable and developed into morphologically normal adults, yet they had decreased fertility. Consistent with the idea that Ski7 participates in remodeling the transcriptome during the oocyte-to-embryo transition, transcriptome profiling identified stage-specific mRNA targets of Ski7. Genes upregulated in *ski7* mutants were generally lowly expressed in wild type, suggesting that Ski7 maintains low transcript levels for this subset of genes. GO enrichment analyses of genes mis-regulated in *ski7* mutants implicated Ski7 in the regulation of redox processes. This was confirmed experimentally by an increased resistance of *ski7* mutant embryos to reductive stress. Overall, our results provide first insights into the physiological role of vertebrate Ski7 as an important post-transcriptional regulator during the oocyte-to-embryo transition.

## Introduction

Post-transcriptional regulation of gene expression plays a major role in defining the amount of RNA and protein in a cell. Apart from ensuring normal cell physiology and maintaining cell fate, post-transcriptional control is important for the speed and robustness of cell fate changes during developmental transitions (Schier, 2007; Tadros & Lipshitz, 2009).

A crucial aspect of post-transcriptional control is RNA decay, which removes unneeded or aberrant RNAs. In eukaryotes, canonical RNA degradation initiates with the shortening of the polyA-tail (Wolf & Passmore, 2014). Following deadenylation, RNAs are degraded by the 5’-to-3’ and/or 3’-to-5’ decay pathways (Houseley & Tollervey, 2009). The 5’-to-3’ degradation pathway is initiated by the removal of the 5’ cap by the Dcp1/Dcp2 decapping complex, followed by transcript elimination by the 5’-to-3’ exonuclease Xrn1 (Sun *et al*, 2013). Alternatively, deadenylated RNAs are degraded in a 3’-to-5’ manner by the RNA exosome complex and several auxiliary cofactors (Lebreton & Séraphin, 2008).

The RNA exosome is an essential and highly conserved multiprotein complex composed of ten core factors. The catalytically inactive subunits of the exosome (members of the RNAse PH protein family) form a hexameric ring, which is capped by a trimeric ring of S1/KH domain-containing proteins (Csl4, Rrp40, Rrp4) (Liu *et al*, 2016; Halbach *et al*, 2013). This catalytically inactive nine-subunit complex associates with one of three exoribonucleases, Rrp6/EXOSC10, Rrp44/DIS3, or DIS3L in the nucleolus, nucleus, or cytosol, respectively (Tomecki *et al*, 2010). Besides its distinct, subcellularly restricted nucleases, the exosome binds auxiliary factors in a compartment-specific manner. In the nucleus, the exosome is activated by the RNA helicase Mtr4 (Weick *et al*, 2018). In contrast, the cytosolic exosome is stimulated by the Ski complex (Araki *et al*, 2001).

The Ski complex is an evolutionary conserved protein complex that participates in canonical 3’-to-5’ RNA decay (van Hoof *et al*, 2000; 2002), RNA surveillance (Horikawa *et al*, 2016; van Hoof *et al*, 2002), viral RNA defense (Toh-E *et al*, 1978; Benard *et al*, 1999), and RNA interference (Orban & Izaurralde, 2005). Consistent with facilitating cytosolic exosome-dependent 3’-to-5’ RNA degradation, studies in yeast have shown that mutants in Ski complex subunits are synthetic lethal with mutants in the 5’-to-3’ degradation pathway (Johnson & Kolodner, 1995). However, the Ski complex, which contains the DExH-box helicase Ski2/SKIV2L, the tetratricopeptide repeat scaffold protein Ski3/TTC37, and two copies of the WD40 repeat protein Ski8/WDR61 (Halbach *et al*, 2013), does not directly interact with the exosome. Association with the exosome complex requires the adaptor protein Ski7, which stably binds to the Csl4 exosome subunit via its P4-like motif, and transiently associates with the Ski complex via the Ski3-Ski8 subunits (Wang *et al*, 2005; Araki *et al*, 2001). In yeast Ski7, the exosome and Ski-complex interacting domains are part of the N-terminus (Kowalinski *et al*, 2016; Araki *et al*, 2001), while the C-terminus consists of a GTPase-like domain involved in RNA quality control (non-stop decay pathway (NSD)) (Horikawa *et al*, 2016; Kowalinski *et al*, 2015).

In contrast to yeast, in which Ski7 is encoded by an independent gene locus (Marshall *et al*, 2013; 2018), the recently identified Ski7 homolog in vertebrates (human HBS1LV3) is encoded by an alternative splice isoform of vertebrate *hbs1l* (Kowalinski *et al*, 2016; Marshall *et al*, 2018; Kalisiak *et al*, 2016). Vertebrate Ski7 shares the Ski and exosome complex interacting domains of yeast Ski7, yet it lacks the GTPase domain (Marshall *et al*, 2018; Brunkard & Baker, 2018; Kowalinski *et al*, 2015). Although a handful of studies have started to characterize vertebrate Ski7’s function *in vitro* (Kowalinski *et al*, 2016; Kalisiak *et al*, 2016), Ski7’s *in vivo* function has not yet been investigated in higher eukaryotes.

Here, we report a functional analysis of the auxiliary 3’-to-5’ degradation factor, Ski7, in zebrafish. We found that *ski7* mRNA peaks during the oocyte-to-embryo transition. Regulation of this transition is fully dependent on post-transcriptional mechanisms including RNA degradation since the mature egg and early zebrafish embryo are transcriptionally silent (Schier, 2007; Tadros & Lipshitz, 2009). Due to Ski7’s implication in RNA degradation and its specific timing of expression, we investigated Ski7’s role during the oocyte-to-embryo transition in zebrafish.

## Results

### Ski7 mRNA is highly expressed during the oocyte-to-embryo transition

Analysis of zebrafish transcriptome (Herberg *et al*, 2018; Pauli *et al*, 2012) and translatome (Pauli *et al*, 2014; Chew *et al*, 2013) data revealed an *hbs1l* splice variant that is highly expressed and translated during early stages of embryogenesis (**Fig. 1A, B**). Conservation analysis of this splice isoform showed that it encodes zebrafish Ski7 (**Fig. 1C**; **Sup. Data 1**). This is consistent with published data from other eukaryotes (Kowalinski *et al*, 2016; Kalisiak *et al*, 2016; Marshall *et al*, 2018). The zebrafish *ski7*-specific exon (exon 5) contains the previously annotated Ski7-like motif and the most conserved region of Ski7, the so-called patch 4-like (P4) motif (Kowalinski *et al*, 2016; Brunkard & Baker, 2018) (**Fig. 1C**). Analysis of *ski7* mRNA levels during oogenesis, in mature eggs, and during embryogenesis showed that *ski7* mRNA peaks during the oocyte-to-embryo transition: while expression of *hbs1l* remains constant, we found that *ski7* mRNA is 13.5-fold higher expressed than *hbs1l* in mature eggs, but 2-fold lower at the beginning of oogenesis and in 5-day old larvae (**Fig. 1B**). A similar trend was observed in mouse (**Sup. Fig. 1A**). Our finding that *ski7* mRNA is enriched in the mature egg raised the possibility that it may be important during the oocyte-to-embryo transition, which is characterized by large-scale changes to RNA and protein content.

**Fig. 1.**
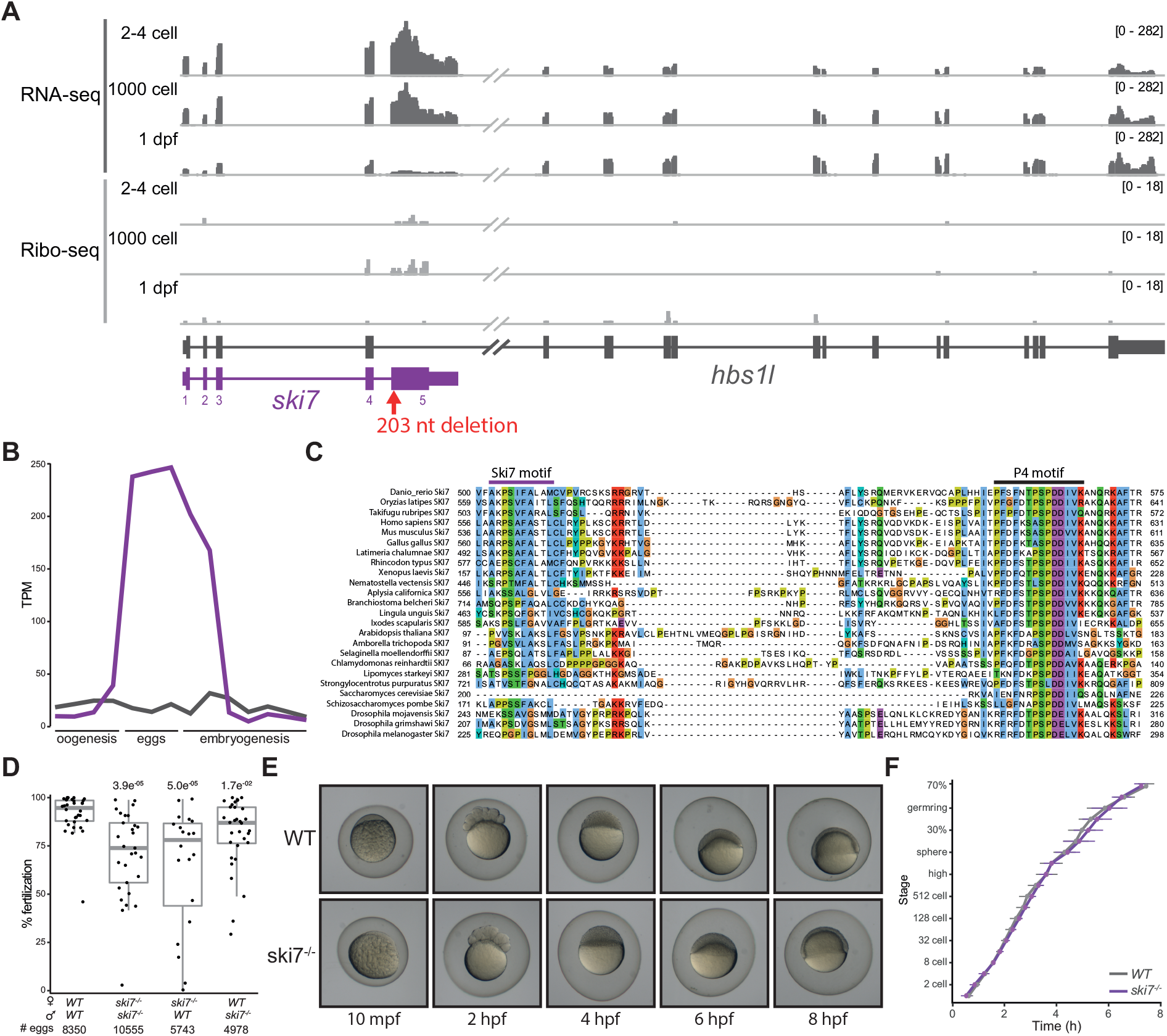
Ski7 is highly expressed during the oocyte-to-embryo transition. A) Expression profile of the *hbs1l* (black)/*ski7* (purple) locus during embryogenesis. *Ski7* is highly expressed in the early embryo. RNA-seq is shown in dark grey and Ribo-seq in light grey; numbers in brackets indicate expression levels; the red arrow indicates the site of the *ski7* mutation. B) *Ski7* mRNA peaks during the oocyte-to-embryo transition. Transcript-specific quantification of *ski7* and *hbs1l* RNA from polyA^+^ RNA-seq during oogenesis, in mature eggs (before and after egg activation, and in fertilized eggs) and during early embryogenesis (from 2-4 cell to 5 dpf (Pauli *et al*, 2012)); TPM, transcripts per million. C) Ski7 protein is highly conserved. Protein sequence alignment of the conserved C-terminus of Ski7 across different organisms. The two most conserved motifs, the Ski7-like motif and the patch 4-like (P4) motif, are highlighted. D) *Ski7^−/−^* fish have reduced fertility. The plot shows fertilisation rates of wild-type (*WT*) and *ski7^−/−^* fish of the indicated crosses. Statistical test: Kruskal-Wallis with Dunn’s multiple comparison test. E) *Ski7^−/−^* embryos develop normally. Representative images of *WT* and *ski7^−/−^* embryos from 10 minutes post-fertilisation (mpf) to 8 hours post-fertilisation (hpf). F) Quantification of embryo development in *WT* and *ski7^−/−^* embryos. *Ski7^−/−^* embryos develop at normal rate.

### Zebrafish Ski7 is not essential for survival

To analyze the function of Ski7 *in vivo* in a vertebrate organism, we generated zebrafish lacking full-length Ski7 protein. To not interfere with Hbs1l expression, we targeted the beginning of the *ski7*-specific exon 5 using CRISPR/Cas9 (**Fig. 1A**; **Sup. Fig. 1B**). We obtained mutant fish carrying a 203-nucleotide deletion resulting in a frameshift mutation after leucine 183 (full-length Ski7: 576 amino acids), which is predicted to generate a truncated Ski7 protein lacking both conserved Ski7-like and P4-like motifs (**Sup. Fig. 1B**). This mutant will hereafter be referred to as *ski7*^−/−^. *Ski7^−/−^* adult fish are viable yet show reduced fertility compared to wild type (Kruskal-Wallis with Dunn’s multiple comparisons test, p = 3.9e^−05^) (**Fig. 1D**). Specifically, reciprocal crosses of *ski7^−/−^* mutant fish with wild-type fish showed a more pronounced subfertility phenotype if Ski7 was lacking in the female as opposed to the male (64.4% and 81.3% average fertilization rates, respectively) (**Fig. 1D**), which suggests a role of maternally provided Ski7. However, *ski7^−/−^* embryos that were successfully fertilized and underwent cell cleavage did not display any morphological defects or developmental delays during embryogenesis (**Fig. 1E, F**) and developed into morphologically normal adult fish. The reduced fertility of *ski7*^−/−^ mutants combined with the lack of an overt developmental phenotype suggest that Ski7’s function is not essential but rather regulatory during the oocyte-to-embryo transition.

### Zebrafish Ski7 interacts with the RNA exosome

Ski7 is known to interact with the cytoplasmic RNA exosome in yeast (van Hoof *et al*, 2000), plants (Brunkard & Baker, 2018), and human cells (Kalisiak *et al*, 2016; Kowalinski *et al*, 2016) by binding to the cap protein Csl4 via multiple interactions, including its highly conserved P4 motif. However, transcriptomic analyses revealed that components of the zebrafish exosome are relatively lowly expressed during the oocyte-to-embryo transition (**Fig. 2A**), leaving it unclear whether zebrafish Ski7 can interact with the exosome during this time frame. To determine whether the interaction of Ski7 with the exosome via the P4 motif was conserved in zebrafish embryos, we conducted co-immunoprecipitation (co-IP) experiments using an *in vitro* synthesized, biotinylated zebrafish Ski7-P4 peptide. Incubation of the peptide with early zebrafish embryo lysate followed by shot-gun mass spectrometry identified 7 of 10 core components of the cytoplasmic exosome as top enriched Ski7-P4-interacting proteins (**Fig. 2B**; **Sup. Data 2**). These results suggest that zebrafish Ski7 is able to interact with the exosome in early embryos via its P4 motif.

**Fig. 2.**
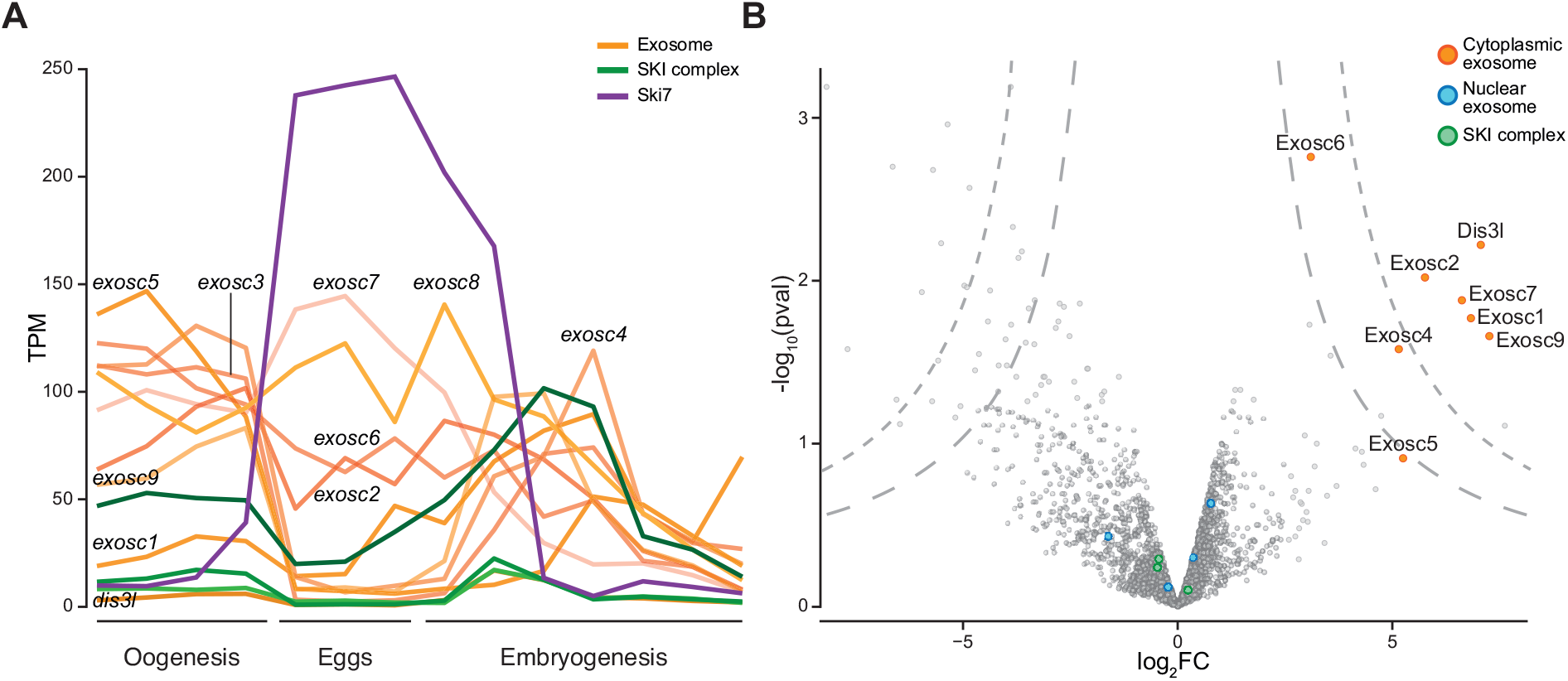
Zebrafish Ski7 interacts with the cytoplasmic exosome. A) mRNA expression levels of *ski7* (purple), subunits of the cytoplasmic RNA exosome (orange) and the SKI complex (green). B) Ski7 associates with the cytoplasmic exosome in early embryos. Volcano plot of Ski7-P4 peptide-interacting proteins from wild-type zebrafish embryo lysates (128-512-cell stage) identified by mass spectrometry. Subunits of the cytoplasmic exosome (orange) are enriched, while nuclear exosome auxiliary factors (blue) and the SKI complex (green) are not enriched.

### Ski7 regulates transcripts in a time-specific manner

As a first step towards characterizing Ski7’s regulatory role during the oocyte-to-embryo transition, we compared the transcriptomes of wild type and *ski7^−/−^* mutants using RNA sequencing. Given our evidence that zebrafish Ski7 can interact with the cytoplasmic exosome (**Fig. 2B**), mRNAs degraded in a Ski7-dependent manner were expected to be stabilized in *ski7*^−/−^ mutants. To identify putative Ski7-dependent mRNA targets, polyA^+^ RNA was isolated at eleven time points spanning the oocyte-to-embryo transition (period 1: oogenesis (O1 = I, O2 = II, O3 = III, O4 = IV & V (Nair *et al*, 2012; Selman *et al*, 1993)), period 2: eggs (INA = inactive, ACT = activated, FER = fertilized), and period 3: embryogenesis (E1 = 2-cell, E2 = 64-cell, E3 = 1000-cell, E4 = sphere) (**Fig. 3A**). Principle component analysis (PCA) of transcript expression levels of wild type and *ski7^−/−^* mutants across this time series revealed that samples clustered primarily by developmental period (oogenesis vs. embryogenesis (PC1)) and time (early vs. late (PC2)) and not by genotype (**Fig. 3B**). Due to the known large-scale gene expression changes between oogenesis and embryogenesis, we suspected that differences between wild-type and *ski7^−/−^* mutant transcriptomes might be masked. To address this, we performed PCA for each of the three periods separately. Notably, this analysis revealed a clear separation by genotype for each period (**Fig. 3C**). This suggests that *ski7*^−/−^ mutants indeed differ in their transcriptomes from wild types during the oocyte-to-embryo transition.

**Fig. 3.**
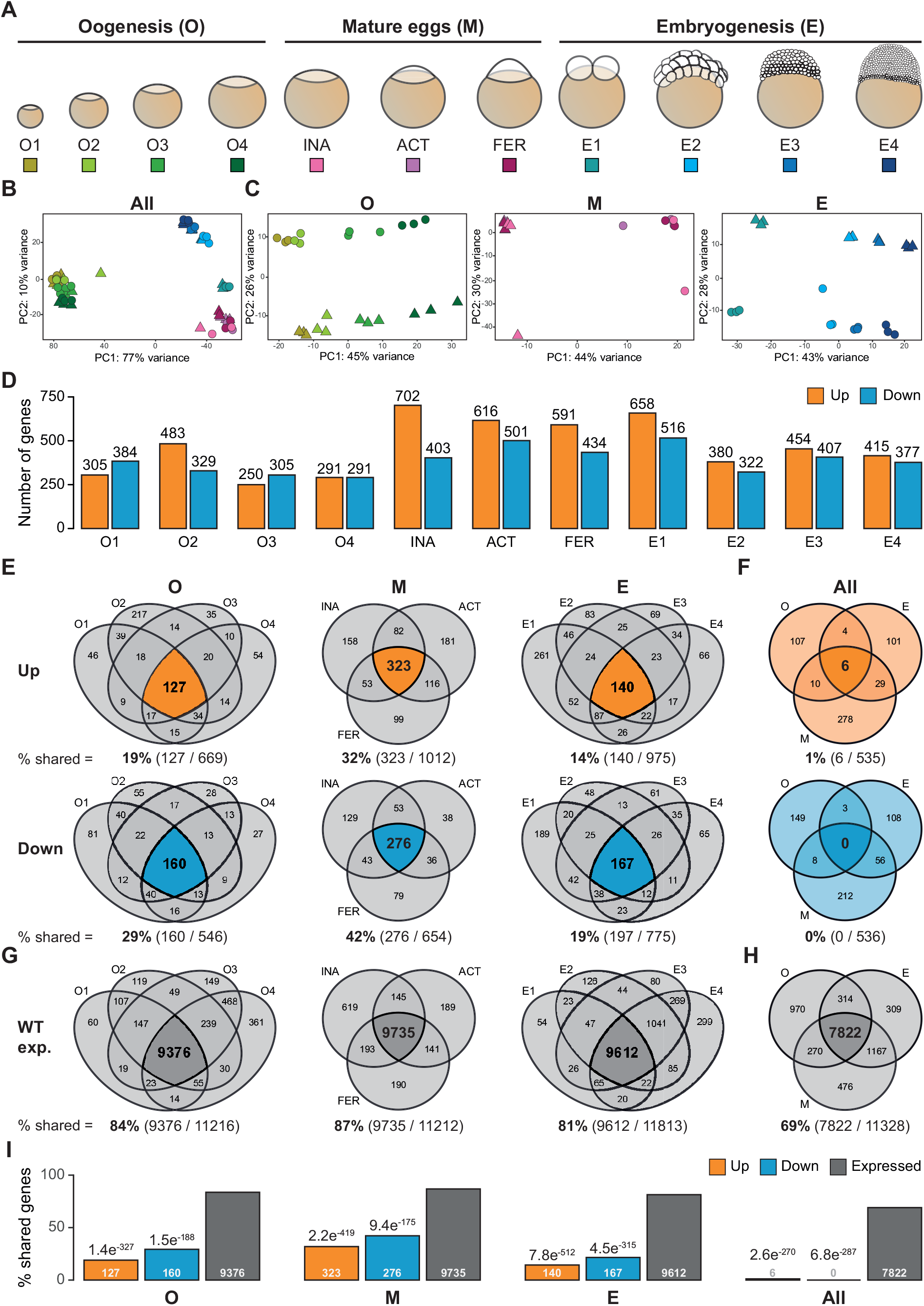
Ski7 modulates the transcriptome during the oocyte-to-embryo transition. A) Schematic representation of the stages used for RNA-seq during three consecutive developmental periods: oogenesis, mature eggs and embryogenesis. B, C) Principal Component Analyses (PCA) of all time points (B) or individual periods (C) used for the RNA-seq comparison of *WT* (circle) and *ski7*^−/−^ (triangle). B) Samples show clustering by developmental time. C) Samples within each period (oogenesis, mature eggs, and embryogenesis) are separated by time and genotype. The colour code corresponds to the different stages from Fig. 2A. D) Differentially expressed genes (DEGs) identified per stage (orange: up-regulated in *ski7*^−/−^ mutants; blue: down-regulated in *ski7*^−/−^ mutants). E) Venn diagrams of DEGs identified in each stage, highlighting shared DEGs per period (orange up-regulated; blue down-regulated). F) Venn diagrams of shared DEGs per period (orange: up-regulated in *ski7*^−/−^ mutants; blue: down-regulated in *ski7*^−/−^ mutants), showing low to no overlap in DEGs between periods. G) Venn diagrams of expressed genes in wild types identified in each stage, highlighting shared genes per period. H) Venn diagram of shared expressed genes in wild type per period, showing high overlap in expressed genes in wild type. I) Percentage of shared DEGs (orange: up-regulated; blue: down-regulated) and expressed genes in wild type (grey) per period (oogenesis, eggs, embryogenesis) and during the full time-course (All). P-values from hypergeometric test from comparison of the number of up- or down-regulated genes versus expressed genes.

Analysis of differentially expressed genes (DEGs) between wild-type and *ski7^−/−^* mutants identified 1215, 1666, and 1750 DEGs (*P*-adjusted ≤ 0.05, ≥ 2-fold change) during oogenesis, in eggs, and during embryogenesis, respectively (**Fig. 3D, E**). Consistent with Ski7’s known role in yeast in 3’-to-5’ mRNA degradation, we found more up- than downregulated genes at each time point throughout the time course: 55% (669/1215), 61% (1012/1666), and 56% (975/1750) of DEGs showed increased expression in *ski7^−/−^* mutants during oogenesis, in eggs, and during embryogenesis, respectively (**Fig. 3D, E**). Comparison of the overlap of identified DEGs between individual time points and between periods revealed that only very few DEGs were shared across all stages (**Fig. 3F**). While there is considerable overlap of DEGs between individual time points within each period (14%-42% overlap) (**Fig. 3E**), DEGs shared between the three periods were rare, making DEGs largely period-specific (1.12% and 0% overlap of up- or down-regulated genes, respectively) (**Fig. 3F)**. This dearth of shared DEGs over time could have resulted from a lack of overlap between transcriptomes at the sampled stages even in a wild-type situation. However, this was not the case since in the wild-type situation, at least 81% of expressed genes were shared within a period (**Fig. 3G**) and 69% among the three periods (**Fig. 3H**). This indicates that expressed genes in wild types overlapped significantly more than the DEGs within and between periods in *ski7^−/−^* mutants (**Fig. 3I**).

What could be possible reasons for the sparsity of shared DEGs in *ski7^−/−^* mutants across the oocyte-to-embryo transition? One possibility is that DEGs are specifically enriched for genes that are exclusively expressed at a specific time point/period in wild types. This would suggest that a common feature of Ski7 targets is time-specific expression. Alternatively, DEGs may be expressed during more than one time point/period in wild types, but only differentially regulated during a defined time in *ski7^−/−^* mutants. This would suggest that the regulation by Ski7 (rather than the presence of its targets) is time-specific. To distinguish between these two possibilities, we assessed the wild-type expression dynamics of each DEG over time. We found that around 90% of Ski7 targets mis-regulated during only one period were expressed during more than one period. In contrast, only around 10% of unique DEGs of each period were restricted in their wild-type expression to that specific period (**Sup. Fig. 2**). This suggests that the vast majority of DEGs arise due to time-specific regulation by Ski7.

### Ski7 targets lowly expressed mRNAs for 3’-to-5’ degradation

Analysis of the expression levels of DEGs in wild types revealed that genes upregulated in *ski7^−/−^* mutants were more lowly expressed in wild types than downregulated and unchanged genes (**Fig. 4A**). This characteristic was consistently observed at each time point during the oocyte-to-embryo transition, which indicates that Ski7 normally targets transcripts with low steady-state levels for RNA degradation. Given that zebrafish Ski7 can interact with the exosome, we hypothesized that genes upregulated in *ski7^−/−^* mutants might have an accumulation of reads at their 3’ ends. To address whether Ski7 targets differ in their read distributions along gene bodies in wild type versus *ski7^−/−^* mutants, we analyzed metagene profiles of up- and downregulated genes as well as unchanged genes for each period. We focused our analyses on the group of shared DEGs within each period (hereafter referred to as ‘shared period DEGs’) as they contain the most confidently identified Ski7-regulated genes per time-window. As expected, metagene profiles from unchanged genes showed no differences between wild types and *ski7^−/−^* mutants (**Fig. 4B**). Additionally, consistent with genes being up- or downregulated in *ski7^−/−^* mutants, *ski7^−/−^* profiles displayed overall higher or lower read coverage compared to wild-type profiles, respectively (**Fig. 4B**). Although this effect was observed across the full transcript, comparison of read densities across individual transcript regions (5’ UTR, coding sequence (CDS), 3’ UTR) revealed a relative enrichment of reads in the CDS and 3’ UTR compared to the 5’ UTR for up-regulated genes (**Sup. Fig. 3**). This bias towards the 3’ end of transcripts supports the idea that zebrafish Ski7 acts similarly to yeast Ski7 in contributing to 3’- to-5’ mRNA degradation.

**Fig. 4.**
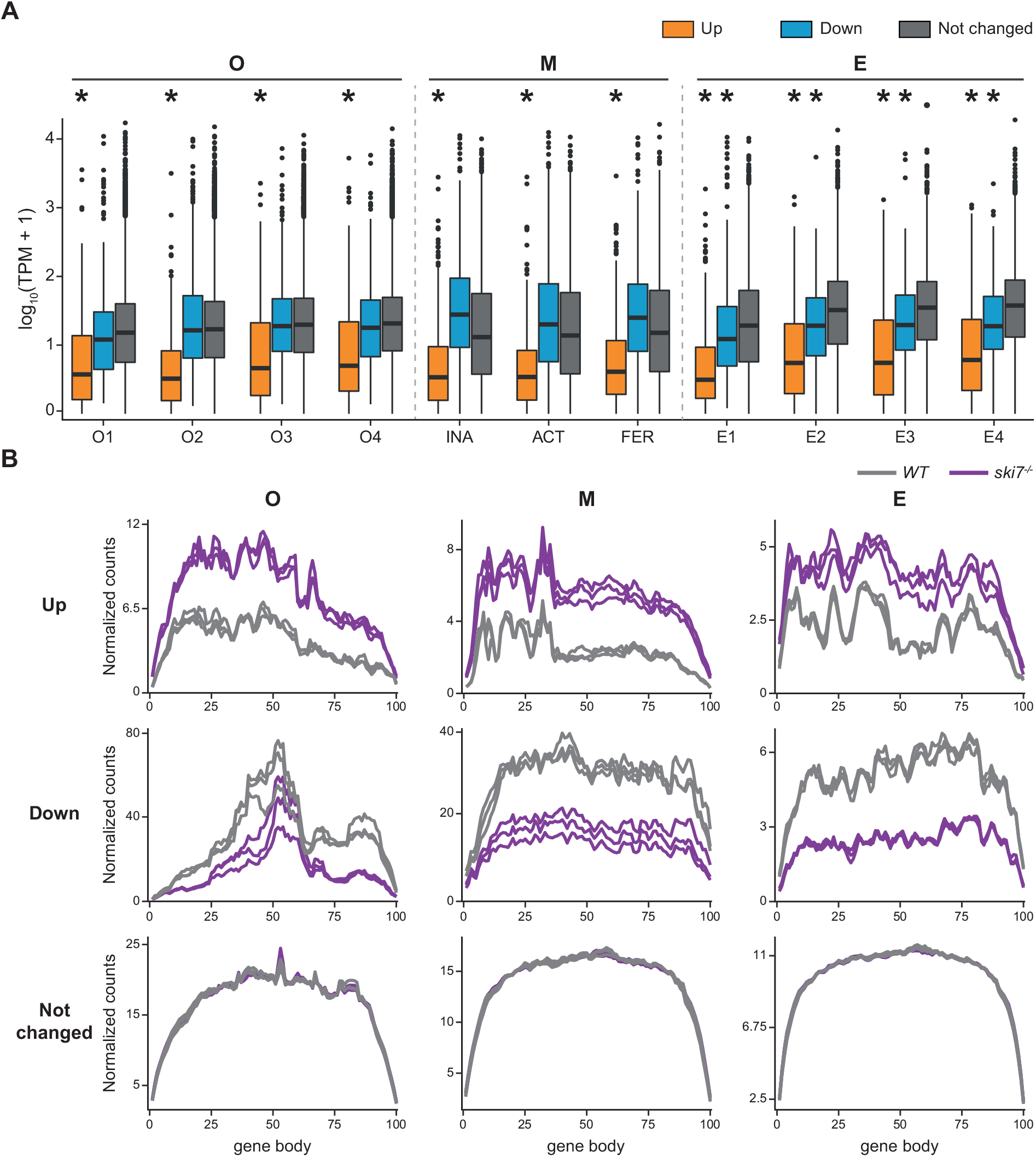
Genes degraded in a Ski7-dependent manner are lowly expressed and degraded from 3’-to-5’. A) Genes up-regulated in the absence of Ski7 show low expression levels in wild type. Expression levels of DEGs in wild type for every stage of the time course, as measured by transcripts per million (TPM). * = p-value < 0.01 obtained by Wilcoxon test. B) Metagene profiles of up-regulated (top), down-regulated (middle), and unchanged genes (bottom) of wild type and *ski7^−/−^* mutants at a representative stage for each period during the oocyte-to-embryo transition (oogenesis = O4, eggs = activated eggs, embryogenesis = E1). *WT*, grey; *ski7^−/−^*, purple.

### Ski7 targets transcripts with diverse features

It has previously been shown that inherent RNA properties, such as transcript and polyA-tail length, codon usage and RNA structure influence RNA stability (Mishima & Tomari, 2016; Bazzini *et al*, 2016; Subtelny *et al*, 2014; Mishima & Tomari, 2017). To investigate whether Ski7 targets share specific transcript features that make them prone for Ski7-dependent regulation, we assessed whether shared period DEGs differ in their intrinsic RNA properties compared to unchanged genes. We observed minor yet significant differences in gene length between the two groups. Genes down-regulated during oogenesis were overall shorter than upregulated or unchanged genes (Kolmogorov-Smirnov p = 4.05e^−6^) (**Sup. Fig. 4A**). This difference could be attributed largely to the shorter lengths of their CDSes and 3’ UTRs (**Sup. Fig. 4B**). In addition, to determine possible differences in the usage of synonymous codons, we calculated the codon adaptation index (CAI) (Carbone *et al*, 2003) of the coding sequences of DEGs and unchanged genes. Genes up-regulated during oogenesis and in eggs showed a lower CAI than down-regulated or unchanged genes (Kolmogorov-Smirnov, p = 0.003 & 0.004) (**Sup. Fig. 5**). Thus, there is an enrichment of rare codons in genes degraded in a Ski7-dependent manner in the female germline.

### Absence of Ski7 confers increased resistance to reductive stress

To determine whether Ski7-dependent mis-regulation of transcripts might result in an imbalance at the protein level, we performed tandem mass tag mass spectrometry (TMT-MS) with wild-type and *ski7^−/−^* embryos at four hours post-fertilization. In total, 76 proteins were found to differ significantly between wild-type and *ski7^−/−^* mutant embryos (33 up-regulated, 43 down-regulated), of which 15 (8 up-regulated, 7 down-regulated) had also been detected as DEGs by RNA-seq (**Sup. Fig. 8A**; **Sup. Data 3**).

We noticed that several proteins upregulated in the absence of Ski7 have been implicated in the regulation of stress response and redox processes (**Sup. Fig. 8B**). To determine whether mis-regulated genes belonged to common pathways or shared biological functions, we performed GO enrichment analyses of DEGs to provide further insights regarding Ski7’s regulatory function during the oocyte-to-embryo transition. Enriched terms in either up- or down-regulated shared period DEGs included processes associated with redox-related processes, cellular respiration, and transporter activity across all three periods (**Fig. 5A**, **Sup. Fig. 6**; for GO analyses per individual stages see **Sup. Fig. 7**).

**Fig. 5.**
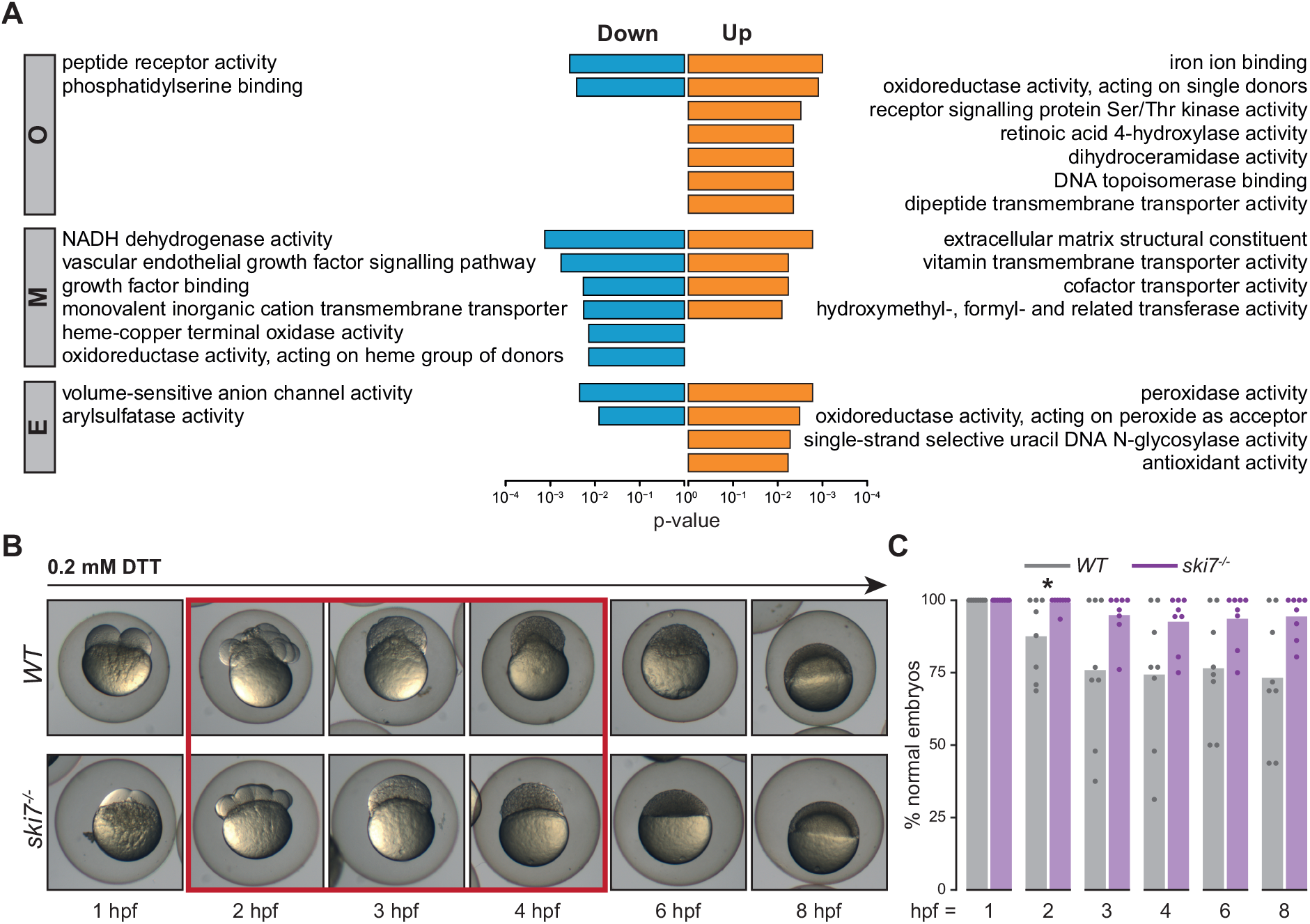
Absence of Ski7 confers increased resistance to reductive stress. A) GO terms enriched in DEGs from *ski7^−/−^* mutants for each period (GO term category ‘Molecular function’). B) *Ski7^−/−^* mutant embryos show increased resistance to DTT-treatment. Representative images of the development of wild-type and *ski7^−/−^* embryos incubated in 0.2 mM DTT. The red box highlights the time-frame during which wild-type embryos show aberrant morphogenesis. C) Quantification of normally developing embryos treated with 0.2 mM DTT during embryogenesis (hpf, hours post-fertilisation). Each dot represents the quantification of an independent experiment (* = p < 0.05). *WT*, grey; *ski7^−/−^*, purple.

Because GO term enrichment analysis and our proteomic analysis indicated a possible link between Ski7 and redox regulation, we assessed whether *ski7^−/−^* embryos differed in their response to redox stress. Treatment of wild-type and *ski7^−/−^* embryos with the reducing agent DTT revealed that *ski7^−/−^* embryos were more resistant to reductive stress than wild-type embryos (**Fig. 5B, C**). This effect was most pronounced between two to four hours post-fertilization though persistent throughout the time course (**Fig. 5B, C**). Overall, these results suggest that embryos lacking Ski7 can cope better with reductive stress, potentially by having elevated levels of redox-regulating factors during the oocyte-to-embryo transition.

## Discussion

In this study, we provide the first functional characterization of vertebrate Ski7 and the consequences of its absence. Zebrafish Ski7, similar to its human and plant homologs (Kalisiak *et al*, 2016; Kowalinski *et al*, 2016; Brunkard & Baker, 2018), is encoded as an alternative splice isoform of *hbs1l*. We found that zebrafish *ski7* but not *hbs1l* mRNA is highly expressed in mature eggs and early embryos, which prompted us to investigate Ski7’s role during the oocyte-to-embryo transition.

Our phenotypic characterization of zebrafish *ski7^−/−^* mutants revealed that Ski7 is not necessary for survival in this vertebrate model system. While this resembles its non-essentiality in yeast (Benard *et al*, 1999), we found that zebrafish *ski7^−/−^* mutants have compromised fertility (**Fig. 1D**), which in a natural environment could result in decreased fitness and ultimately lead to impaired propagation and maintenance of the species. The compromised fertility of *ski7^−/−^* mutants, together with our finding that, once fertilized, *ski7^−/−^* mutant embryos have no further apparent defects in development, suggests that Ski7 mainly acts during oogenesis and/or in the mature egg.

What could Ski7’s function be during the oocyte-to-embryo transition? We have three lines of evidence that zebrafish Ski7 acts – in analogy to the situation in yeast (van Hoof *et al*, 2000; Araki *et al*, 2001) and human cells (Kowalinski *et al*, 2016; Kalisiak *et al*, 2016) - in conjunction with the cytoplasmic RNA exosome in promoting 3’-to-5’ RNA decay: Firstly, we find that zebrafish Ski7 can interact with the RNA exosome (**Fig. 2B**). Secondly, there are more up- than down-regulated genes in the absence of Ski7 (**Fig. 3D**). And lastly, up-regulated genes show an accumulation of reads towards their 3’ ends (**Fig. 4B**; **Sup. Fig. 3**). Our conclusion that the Ski7/RNA exosome complex acts during zebrafish egg development is consistent with the recently reported role for the exosome core component EXOSC10 in the maturation of mouse eggs (Wu & Dean, 2020). While the phenotypic consequences of loss of Ski7 in mice remain to be determined, our analysis of publicly available RNAseq data (Hendrickson *et al*, 2017; Yu *et al*, 2016) revealed that mouse *Ski7* mRNA is also highly expressed in mature eggs and decreases during embryogenesis while *Hbs1l* remains stable. Taken together, Ski7’s function in fine-tuning transcript levels during egg maturation by facilitating 3’-to-5’ RNA degradation might be conserved in vertebrates.

Our data reveal that Ski7 preferentially targets transcripts for degradation whose steady-state levels are low. This suggests possible roles for Ski7 in maintaining low transcript abundances for certain genes and/or reducing transcriptional noise, possibly in conjunction with other mechanisms including the preferential use of rare codons. While the majority of mis-regulated genes in *ski7* mutants are up-regulated, many genes were also found to be down-regulated (**Fig. 3D**). Down-regulated genes could either be due to secondary effects caused by the lack of down-regulation of primary Ski7/RNA exosome target genes, or due to so-far unknown activities of Ski7. This raises the question whether Ski7 acts solely as auxiliary factor of the RNA exosome, or might have additional, non-canonical functions. While we have currently no evidence for alternative functions, we cannot rule out the possibility that Ski7 might interact with other factors in addition to the RNA exosome, particularly in light of the comparatively low mRNA expression levels of the core exosome components during the oocyte-to-embryo transition (**Fig. 2A**). However, our finding that a Ski7-P4 peptide fragment can stably bind to the zebrafish exosome complex in early embryo lysates demonstrates that zebrafish Ski7 can stably interact with the RNA exosome (**Fig. 2B**). This experiment also demonstrates that the zebrafish Ski7-P4 motif (PFSFNTPSPDDIVK) is sufficient to pull down the exosome in zebrafish embryos. This is in contrast to data from human cells in which the Ski7-like motif but not the P4 motif was shown to be sufficient for mediating this interaction (Kalisiak *et al*, 2016). Thus, our data demonstrates a direct interaction between Ski7 and the RNA exosome. Another question that will warrant further investigations is the underlying reason for the relatively small number of shared DEGs among different time frames. Along these lines, several recent studies have challenged the view that the general RNA decay machinery is unspecific. For example, Dcp2 (5’ decapping) and CCR4NOT (3’ deadenylation) target specific subsets of mRNAs depending on inherent transcript features during zebrafish embryo development (Mishima *et al*, 2006; Horvat *et al*, 2018; Fujino *et al*, 2018; Sohrabi-Jahromi *et al*, 2019; Sorenson *et al*, 2018; He *et al*, 2018). However, it remains largely unclear to what extent general cytoplasmic RNA degradation machineries or specific components thereof contribute to the large-scale transcriptome remodeling during development. Our analyses indicate that the lack of shared DEGs among periods is not due to time-restricted expression of the target mRNAs in the wild-type situation but rather time-specific regulation by Ski7. How Ski7’s time-specificity is achieved is still unclear. Possible mechanisms include temporal stabilization of transcripts by RNA-binding proteins or sequestration from the Ski7/RNA exosome complex, as well as yet-to-be-identified temporal Ski7-specific regulators.

Our observation that *ski7^−/−^* mutant embryos are less sensitive to reductive stress was unexpected since previous studies in yeast have indicated the opposite effect: In yeast, Ski7-mediated non-stop decay (NSD) was shown to be required for oxidant tolerance in combination with the Dom34-Hbs1 complex (Jamar *et al*, 2017). In mammals, it is well established that redox regulation, e.g. by regulating the levels of NADPH/NADH, is particularly important during oogenesis and in the egg (Bertoldo *et al*, 2020). De-regulation of NADH/cellular respiration in *ski7^−/−^* eggs could therefore contribute to the reduced fertility, in addition to the mis-regulation of reproductive genes themselves. Consistent with this idea, enriched GO terms for up-regulated genes were associated with regulation of fertilization (egg coat, acrosin binding), while enriched GO terms for downregulated genes included NADH dehydrogenase, oxidoreductase activity, cellular respiration, and the unfolded protein response (**Sup. Fig. 6, 7**). Yet how can we explain the increased tolerance of *ski7^−/−^* mutants to the reductive agent DTT during embryogenesis? In the simplest scenario, Ski7 might negatively regulate redox stress tolerance in zebrafish embryos by fine-tuning the levels of embryonic redox regulators. However, the contrasting observations, namely that *ski7^−/−^* mutant eggs have reduced fertility, but that mutant embryos are resistant to reductive stress, are difficult to explain by a static scenario. We therefore propose an alternative possibility, namely that Ski7 might contribute to the dynamic regulation of redox balance during the oocyte-to-embryo transition. In support of this idea, we found that thioredoxins, cellular respiration and mitochondrial processes are mis-regulated throughout the oocyte-to-embryo transition in *ski7^−/−^* mutants, which could result in a time-dependent imbalance in redox regulation. Evidence for increased oxidative levels after fertilization come for example from *Xenopus*, in which fertilization enhances the production of mitochondrial ROS (Han *et al*, 2018), *C. elegans* (Knoefler *et al*, 2012) and zebrafish embryos (Gauron *et al*, 2016). Additionally, in support of the idea of dynamic redox regulation by Ski7, it was reported that *Drosophila* oocytes lacking an oocyte-specific thioredoxin had increased levels of H_2_O_2_ and were more resistant to DTT than their wild-type siblings (Petrova *et al*, 2018). Apart from the idea that Ski7 could regulate the dynamic redox balance throughout the oocyte-to-embryo transition, another, non-contradicting possibility is that the increased resistance to reductive agents during embryogenesis might be a direct or indirect consequence of compromised egg maturation in the absence of Ski7. As such, counter-measures of the egg combatting the impaired regulation of RNA levels during oogenesis might make the egg better prepared for later encounters of redox stresses. As such, up- regulation of redox-regulators and/or chaperones during oogenesis could enable the embryo to deal better with reductive agents. This is in line with previous findings that mammalian oocytes treated with H_2_O_2_ produced more blastocysts under *in vitro* conditions (Vandaele *et al*, 2010). While we currently don’t know the extent or direct impact of Ski7-mediated regulation on the redox balance during the oocyte-to-embryo transition, one intriguing speculation is that Ski7 production itself might be redox-controlled. Splicing factors have been shown to be differentially regulated by redox stress (Berner *et al*, 2017), opening the possibility that the alternative splicing reaction responsible for generating the Ski7-specific splicing event might at least in part be triggered by the redox-conditions encountered during oogenesis.

Overall, our data supports the idea that Ski7 fine-tunes the 3’-to-5’ RNA degradation machinery, namely the RNA exosome, to aid in balancing redox regulation during the oocyte-to-embryo transition.

## Materials and Methods

### Fish husbandry

Zebrafish (*Danio rerio*) were raised according to standard protocols (28°C water temperature; 14/10 hour light/dark cycle). TLAB fish, generated by crossing zebrafish AB and the natural variant TL (Tupfel Longfin) stocks, served as wild-type zebrafish for all experiments. *Ski7*^−/−^ mutant lines were generated in the TLAB background and are described below. All fish experiments were conducted according to Austrian and European guidelines for animal research and approved by local Austrian authorities (animal protocol GZ: 342445/2016/12).

### Conservation analysis

The last 228 nucleotides (76 amino acids) of the zebrafish Ski7-specific exon (exon 5) were used to perform the conservation analyses. This region included the predicted Ski7-like and P4-like motifs. The zebrafish sequence was taken as a query for iterative PSI-BLAST searches against the NCBI non-redundant protein data base. In the first search round, not only vertebrate orthologs but also sequences from the coral *Stylophora pistillata* and the brachiopod *Lingula unguis* were detected (E-values < 0.001). Further iterations extended to the plant (e.g. *Ananas comosus*, round 2, E-value 2.17e-05) and fungi kingdom (e.g. *Botryotinia calthae*, round 3, E-value 3.73e-09). Significant hits covered both motifs. However, sequences with a weak Ski7-like motif, such as *Drosophila melanogaster* or *Saccharomyces cerevisiae* could not be identified with zebrafish Ski7, but were found with iterative PSI-BLAST searches of related insect or fungi orthologs. Sequences that represented a wide taxonomic range were selected, aligned using mafft (version 7.427 (Katoh *et al*, 2002)) and visualized in Jalview (Waterhouse *et al*, 2009).

### Generation of *ski7^−/−^* mutant fish

Zebrafish *ski7^−/−^* mutants were generated by CRISPR/Cas9-mediated mutagenesis. CHOPCHOP (Montague *et al*, 2014; Labun *et al*, 2019) was used to predict three guide RNAs targeting the *ski7*-specific exon 5:

**gRNA 1:** GCACTGCTAACAGATCACTGAGG
**gRNA 2:** GCCAAACTTCCAAAACCAGGGGG
**gRNA 3:** AGACCAGTGGAGGGAACTAGAGG

T7-promoter sequence- and 3’ overhang-containing DNA oligos were synthesized (Sigma-Aldrich), annealed to a common tracer oligo (AAAAGCACCGACTCGGTGCCACTTTTTCAAGTTGATAACGGACTAGCCTTATTTTAACTTGCTATTTCTAGCTCTAAAAC), and *in vitro* transcribed by T7 polymerase (Ambion MEGAscript T7 transcription kit) according to published procedures (Gagnon *et al*, 2014). In-house produced Cas9 protein and pooled gRNAs were co-injected in the cell of one-cell TLAB zebrafish embryos. Potential founder fish were outcrossed to TLAB wild-type fish. Progeny were screened for carrying a mutation in *ski7* by PCR based on a size difference of the PCR product compared to the wild-type PCR product, using the primers ski7_F: GACTGTATCCAGTGCACATTCA and ski7_R: TTAAAAGAGCCAAGAGGACTGG. Embryos with potential mutations were raised. Adult fish were genotyped by standard fin-clipping procedures. Sanger sequencing of the PCR fragment identified a deletion of 214 nucleotides and an insertion of 11 nucleotides in *ski7’s* terminal exon 5, resulting in a net deletion of 203 nucleotides that causes a frameshift mutation. Heterozygous fish were incrossed to generate homozygous *ski7*^−/−^ mutant fish. The PCR products were detected by 2% gel electrophoresis (WT = 330 nucleotides, *ski7^−/−^* = 127 nucleotides).

### Quantification of fertilization rates

To quantify fertilization rates, individual male and female fish were set up together in breeding cages the night before mating. The next morning, fish were put together by removing the divider and allowed fish to mate for approximately 30 minutes. Eggs were collected from individual tanks and kept at 28°C in blue water (E3 medium with 0.1% methylene blue per liter of medium). Fertilization rates were determined between 5 and 6 hours post-fertilization.

### Phenotypic analyses and imaging

Quantification and staging of embryos were performed by live cell imaging on a Celldiscoverer 7 automated microscope (Zeiss). Wild-type and *ski7^−/−^* embryos were dechorionated with pronase (1 mg/ml). Dechorionated embryos were mounted in 0.5% low melt agarose within the first 30 minutes post-fertilization. Mounted embryos were kept in six-well plates in E3 medium with 0.1% methylene blue per liter of medium at 28°C during imaging. Images were adquired every 5 minutes during the first 7 hours. Quantification and staging during the first hour post-fertilization was performed manually every five minutes in the fish facility (28°C) under a stereo microscope.

### Ski7-P4 peptide pull-down and MS analysis

The zebrafish Ski7-P4 peptide (HHIEPFSFNTPSPDDIVKANQRK) was synthesized *in vitro* and N-terminally biotinylated (Biotin-Ahx-HHIEPFSFNTPSPDDIVKANQRK). The peptide contained the homologous region of yeast Ski7 that mediates the interaction with the Csl4 exosome protein (second half of helix 3 and first half of helix 4 in yeast) (Kowalinski *et al*, 2016). To prepare zebrafish embryo lysates, embryos were dechorinated with pronase (1 mg/ml). About 250 embryo caps per sample (each experiment was performed in triplicates) were manually dissected from 128-cell to 512-cell stage wild-type zebrafish embryos and immediately frozen in liquid nitrogen. Embryo caps were homogenized with a plastic pestle in 1 ml of cold binding buffer (150 mM KaOAc, 300 μl Hepes-NaOH pH 7.4, 5 mM MgSO_4_, 0.1% NP-40, 5mM DTT, complete EDTA-free protease-inhibitors) and cleared by 1 minute centrifugation at 15000 rpm at 4°C. The biotin pull-down was performed according to a protocol adapted from (Nishida *et al*, 2009). In brief, 30 μl of Streptavidin Dynabeads (MyOne, Invitrogen) per sample were washed with PBS and binding buffer. Beads were incubated with 2 μg of biotinylated Ski7-P4 peptide in 500 μl of cold binding buffer (2 hours at 4°C). Three samples of beads not incubated with biotinylated peptide served as negative controls for the pull-down. After washing of the beads (five times with 500 μl of binding buffer; total wash time 2 hours at 4°C), beads were incubated with 1 ml embryo lysate for 2 hours at 4°C on a rotating wheel. Beads were placed on a magnetic stand and washed six times with 1 ml of cold binding buffer (total wash time 2-4 hours at 4°C). After washing, bound proteins were eluted by adding 30 μl of hot (95°C) 2x SDS-PAGE sample buffer to the beads. For mass-spec analysis of the eluates, samples were run into a SDS-PAGE gel for a short time (without allowing separation of the proteins by size) and stained by Coomassie Blue. Gel elution of the Coomassie-stained band and tryptic digest followed standard procedures.

For LC–MS/MS, digested peptides were separated using a Dionex UltiMate 3000 HPLC RSLC nano system (Thermo Fisher Scientific) coupled to a Q Exactive mass spectrometer (Thermo Fisher Scientific), equipped with a Proxeon nanospray source (Thermo Fisher Scientific). Peptides were loaded onto a trap column (PepMap C18, 5 mm × 300 μm ID, 5 μm particles, 100 Å, Thermo Fisher Scientific) at a flow rate of 25 μL min^−1^ using 0.1% TFA as mobile phase. After 10 min, the trap column was switched in line with the analytical column (PepMap C18, 500 mm × 75 μm ID, 2 μm, 100 Å, Thermo Fisher Scientific). A 105/165/225 min gradient from buffer A (water/formic acid, 99.9/0.1, v/v) to B (water/acetonitrile/formic acid, 19.92/80/0.08, v/v/v) was applied to elute the peptides at a flow rate of 230 nl min^−1^. The Q Exactive HF mass spectrometer was operated in data-dependent mode, using a full scan (m/z range 380-1500, nominal resolution of 60,000, target value 1E6) followed by MS/MS scans of the 10 most abundant ions. MS/MS spectra were acquired using normalized collision energy of 27%, isolation width of 1.4 m/z, resolution of 30.000 and the target value was set to 1E5. Precursor ions selected for fragmentation (excluded charge state 1, 7, 8, >8) were put on a dynamic exclusion list for 20/40/60 s, depending on the gradient length. Additionally, the minimum AGC target was set to 5E3 and the intensity threshold was calculated to be 4.8E4. The peptide match feature was set to preferred and the exclude isotopes feature was enabled. For peptide identification, the RAW files were loaded into Proteome Discoverer (v2.1.0.81, Thermo Scientific). All MS/MS spectra were searched using MSAmanda v2.1.5.8715 (Dorfer *et al*, 2014) against a custom *Danio rerio* protein database (based on GRCz10 and CRC64 non-redundant Ensembl 86 release: 42,188 sequences, 22,758,666 residues). The following search parameters were used: fixed modification: beta-methylthiolation on cysteine; variable modifications: oxidation on methionine, phosphorylation on serine, threonine and tyrosine, deamidation on asparagine and glutamine, acetylation on lysine, methylation on lysine and arginine, di-methylation on lysine and arginine, tri-methylation on lysine, ubiquitinylation on lysine, glycosylation (HexNac) on asparagine, serine and threonine as well as glutamine-to-pyro-glutamic-acid-transformation on peptide-N-terminal glutamine. Monoisotopic masses were searched within unrestricted protein masses for tryptic enzymatic specificity. The peptide mass tolerance was set to ±5 ppm and the fragment mass tolerance to ±0.03 Da. The maximal number of missed cleavages was set to two. The result was filtered to 1% FDR on peptide level using Percolator algorithm integrated in Thermo Proteome Discoverer. The localization of the modification sites within the peptides was performed with the tool ptmRS, which is based on the tool phosphoRS (Taus *et al*, 2011). Label-free quantification of peptides was performed using the in-house developed tool apQuant (Doblmann *et al*, 2019). Proteins were quantified by summing up all peptides before performing subsequent iBAQ normalization (Schwanhäusser *et al*, 2011). Differentially expressed proteins were determined using limma (Smyth, 2005). P-values were adjusted for multiple testing as implemented in the limma R package (Ritchie *et al*, 2015).

### RNA isolation and sequencing

Total RNA was extracted using the standard TRIzol (Invitrogen) protocol. Samples from individual stages of three different periods (oogenesis, mature eggs, embryogenesis) were collected as follows: To obtain different stages of oogenesis, ovaries were dissected from three different females from each genotype. Oocytes were mannualy sorted based on differences in size (O1 – smallest size; O4 – largest size but smaller than mature eggs) and kept in oocyte sorting medium (Leibovitz’s medium, plus 0.5% BSA, pH 9.0 adjusted with 5 M NaOH (Nair *et al*, 2012)). To obtain different stages of mature eggs, inactive, active and fertilized eggs were collected from the same female fish. Three independent biological replicates were collected of each stage (three different females, between 35 to 60 eggs per condition). Briefly, single female fish were put together with male fish for standard mating. Eggs that were laid within the first minute after putting them together were collected at 10 minutes post-fertilization and homogenized in TRIzol (fertilized eggs). Fertilization of the eggs was quantified after 3 hours and only samples in which the rate of fertilization was higher than 60% were used. The females used for collecting fertilized eggs were immediately anesthetized using 0.1% Tricaine and subjected to squeezing to collect the remaining two stages, inactive and activated eggs. Half of the squeezed eggs were immediately homogenized in TRIzol (inactive eggs). The remaining half of the eggs was activated by adding fish water (E3 medium with 0.1% methylene blue per liter of medium) and incubated for 10 minutes before collection and homogenization in TRIzol (activated eggs). To obtain different stages during embryogenesis, between 20 to 30 embryos per time point were collected at the indicated stages and homogenized in TRIzol. Total RNA was isolated and assessed for quality and quantity based on analysis on the Fragment Analyzer. 2 μg of total RNA was used per sample as input for PolyA+ RNA selection with the poly(A) RNA Selection Kit (LEXOGEN). Stranded cDNA libraries were generated using NEBNext Ultra Directional RNA Library Prep Kit for Illumina (New England Biolabs) and indexed with NEBNext Multiplex Oligos for Illumina (Dual Index Primer Set I) (New England Biolabs). Library quality was checked on the Fragment Analyzer and sequenced on a Illumina Hiseq 2500 on SR100 mode.

### RNA-seq data processing

RNA-seq reads were processed according to standard bioinformatic procedures. First, BAM files were converted to fastq files using samtools (v1.9) (Li *et al*, 2009). Barcoded libraries were demultiplexed using fastx-toolkit (v0.0.14) (http://hannonlab.cshl.edu/fastx_toolkit/) and customized perl scripts. Sequencing adapters were trimmed with bbduk command from the BBmap package (v38.26) (Bushnell, 2014) and only reads longer than 20 bases were kept for downstream analyses. Filtered reads were mapped to Ensembl 92 gene models (downloaded 2018.08.01) and the GRCz11 *Danio rerio* genome assembly, with Hisat2 v2.1.0 (Kim *et al*, 2019) (--rna-strandness R -k 12 –no-unal). Libraries that had fewer than 0.5 million reads were not considered for the analyses; thus, for wild-type inactive eggs and wild-type fertilized eggs only two replicates were included for the analysis. Quantification at the gene level (transcript per million (TPM)) was perfomed using Kallisto (v0.43.0) (Bray *et al*, 2016) and customized perl scripts.

### Differential gene expression analyses

HTSeq (v0.9.1) (Anders *et al*, 2015) was employed to count the uniquely mapped reads. Differential gene expression was performed on raw counts using DESeq2 (v1.22.2) (Love *et al*, 2014) with default parameters and size factors estimated from global normalization. The three different periods were analyzed separately, except for the generation of the combined PCA plot (Fig. 2B). Only genes that had cpm (counts per million) values equivalent to at least 5 raw counts in at least two libraries per period were considered for the analyses. Genes were considered as differentially expressed if they had an adjusted p-value < 0.05 and a log_2_ fold change ≤ −1 or ≥ 1. Principal component analyses were performed after a regularized log transformation using the 1000 most variable genes.

### Analyses of RNA features

For all analyses, only the longest isoform was used for genes with multiple isoforms. For metagene profiles, full length transcripts of at least 100 nucleotides were used. BAM files containing mapped reads to the transcriptome were converted to BedGraph files using Bedtools (v2.27.1) (Quinlan & Hall, 2010) and plotted using customized R scripts. To analyse the lengths of individual gene body regions (5’ UTRs, CDSs and 3’ UTRs), the Ensembl 92 annotation file was used. Read density by region was defined as number of reads per length of the region of interest. Number of reads per region was calculated using HTSeq (v0.9.1) (Anders *et al*, 2015) and feature type (−f) as ‘five_prime_UTR’, ‘CDS’, or ‘three_prime_UTR’. If a read belonged to two distinct regions, it was counted for both of them. The ratio of the read densities (read density in *ski7* mutants versus read density in WT) was used to assess relative enrichments of reads over distinct gene body regions. If wild-type and *ski7* read densities were zero, the value was not considered for the calculation of the ratio. Codon adaptation index (CAI) of the coding sequences of annotated genes was calculated using a command line version of the CAIcal program (v1.4) (Puigbò *et al*, 2008) (zebrafish codon usage table obtained from the codon usage database (https://www.kazusa.or.jp/codon/)) and standard genetic code (−g 1). The CAI was calculate per gene (−s y). As control, the CAI was calculated for a number of random sequences (−n) equal to the size of the group being compared (or 500 if the group was bigger than 1000 sequences) and with the number of codons (−l) equal to the average length of the CDS being analyzed. GO term enrichment analyses were performed for up- and down-regulated genes separately using topGO (v2.34.0) (Alexa & Rahnenfuhrer, 2019). A reduction of terms for visualization was performed with the online tool REVIGO.

### TMT-MS

TMT-MS was performed in biological triplicates. For each sample, 200 embryos from individual parents were collected at 4 hours post-fertilization. Embryos were dechorionated (1 mg/ml pronase) and batch-deyolked following (Link *et al*, 2006). Samples were lysed by sonication in SDT buffer (4% SDS, 0.1 M DTT, 0.1 M Tris-HCl pH 7.5). For downstream normalization, the STD buffer was supplemented with 1 pmol of dCas9 protein and Lambda exonuclease protein per 10 μl buffer. Samples were diluted with 200 μl of 8 M urea, 0.1 M Tris-HCl, pH 8.5 and centrifuged at 14,000 g for 20 minutes using a 30 kDa mass-cutoff 0.5 ml Amicon centrifugal filter (Merck). For alkylation of cysteines, samples were incubated for 30 minutes in the dark after addition of 100 μl of 0.1 M IAA, 8 M urea, 0.1 M Tris-HCl pH 8.5 and then centrifuged at 14000 g for 20 minutes. Samples were washed with 100ul of 8 M urea, 0.1 M Tris-HCl pH 8.5 and 100 μl of 100 mM Hepes pH 7.6 by centrifugation at 14,000 g for 20 min (four wash steps in total). Tryptic digests were performed overnight after addition of 46 μl of 100 mM Hepes pH 7.6 and 4 μl of 1 μg/μl Trypsin Gold (Promega).

TMT Labelling: Samples were acidified with 10% TFA and cleaned up with 50 mg Sep-Pak C18 columns (Waters), freeze-dried overnight and then dissolved in 100 μl of 100 mM Hepes. Samples were run on a monolithic column for quantification to use not more than 100 μg per sample for TMT labeling. Samples were labelled with TMT according to manufacter’s instructions (Thermo Fisher). Labelled samples were pooled and cleaned up via a 50 mg Sep-Pak C18 column (Waters) and separated into fractions using a SCX system (UltiMate system (Thermo Fisher) and TSKgel SP-25W column (Tosoh Bioscience; 5 μm particles, 1 mm ID × 300 mm), flow rate of 30 μl/min). For the separation, a ternary gradient was used: buffer A (5 mM phosphate buffer pH 2.7, 15% ACN), buffer B (5 mM phosphate buffer pH 2.7, 1 M NaCl, 15% ACN), and buffer C (5 mM phosphate buffer pH 6, 15% ACN). 60 fractions were collected and stored at −80°. The fractions were analysed with LC-MS. The nano HPLC system used was an UltiMate 3000 HPLC RSLC nano system (Thermo Scientific) coupled to a Q Exactive HF-X mass spectrometer (Thermo Scientific), equipped with a Proxeon nanospray source (Thermo Scientific). Peptides were loaded onto a trap column (Thermo Scientific, PepMap C18, 5 mm × 300 μm ID, 5 μm particles, 100 Å) at a flow rate of 25 μL min^−1^ using 0.1% TFA as mobile phase. After 10 min, the trap column was switched in line with the analytical column (Thermo Scientific, PepMap C18, 500 mm × 75 μm ID, 2 μm, 100 Å). A 180 min binary gradient from buffer A (water/formic acid, 99.9/0.1, v/v) to B (water/acetonitrile/formic acid, 19.92/80/0.08, v/v/v) was applied to elute the peptides at a flow rate of 230 nl min^−1^. The Q Exactive HF-X mass spectrometer was operated in data-dependent mode, using a full scan (m/z range 375-1650, nominal resolution of 120,000, target value 3E6) followed by MS/MS scans of the 10 most abundant ions. MS/MS spectra were acquired using normalized collision energy of 35, isolation width of 0.7 m/z, resolution of 45.000, a target value of 1E5, and maximum fill time of 250 ms. For the detection of the TMT reporter ions, a fixed first mass of 110 m/z was set for the MS/MS scans. Precursor ions selected for fragmentation (exclude charge state 1, 7, 8, >8) were put on a dynamic exclusion list for 60 s. Additionally, the minimum AGC target was set to 1E4 and intensity threshold was calculated to be 4E4. The peptide match feature was set to preferred and the exclude isotopes feature was enabled.

For peptide identification, the RAW files were processed in Proteome Discoverer (v2.3.0.523, Thermo Scientific). All MS/MS spectra were searched using MSAmanda (v2.0.0.141114) (Dorfer *et al*, 2014) against a custom *Danio rerio* protein database (based on GRCz11 and annotated gene models from the Ensembl 92 release: 38,422 sequences, 21,882,367 residues supplemented with the sequences of the spike-ins). The parameters used were as follows: fixed modification: Iodoacetamide derivative on cysteine; variable modifications: deamidation on asparagine and glutamine, oxidation on methionine, phosphorylation on serine, threonine and tyrosine, sixplex tandem mass tag on lysine, as well as carbamylation and simplex tandem mass tag on peptide N-termini. Trypsin was defined as the proteolytic enzyme allowing for up to two missed cleavages. Monoisotopic masses were searched within unrestricted protein masses for tryptic enzymatic specificity. The peptide mass tolerance was set to ± 5 ppm and the fragment mass tolerance to ± 15 ppm. The maximal number of missed clavages was set to two. Identified spectra were FDR-filtered to 1.0 % on peptide level using the Percolator algorithm integrated in Thermo Proteome Discoverer. The localization of the modification sites within the peptides was performed with ptmRS, which is based on phosphoRS (Taus *et al*, 2011). Peptides were quantified based on Reporter Ion intensities as extracted by the “Reporter Ions Quantifier”-node as implemented in Proteome Discoverer. Proteins were quantified by summing up unique and razor peptides. Protein area normalization was done using the spiked-in proteins dCas9 and Lambda exonuclease. Differentially expressed proteins were determined using limma (Smyth, 2005) and p-values were adjusted for multiple testing using the limma R package (Ritchie *et al*, 2015).

### Reductive agent sensitivity test in zebrafish embryos

To assess the sensitivity of wild-type and mutant embryos to DTT, embryos were transferred to 0.2 mM DTT in blue water (E3 medium with 0.1% methylene blue per liter of medium) at 30 minutes post-fertilization. Embryos were kept in this medium for the duration of the experiment. Untreated embryos (not incubated in DTT) were kept alongside as controls. Manual examination of defects was performed at every hour for the first 8 hours post-fertilization.

### Statistical analyses

Data visualization and all statistical analyses were performed using R (v3.5.1).

## Acknowledgements

We thank the proteinchemistry facility at the ViennaBiocenter, particularly Richard Imre, Michael Schutzbier and Gerhard Dürnberger, for TMT-MS sample preparation and support in proteomics data analyses; Karin Aumayer and her team of the biooptics facility at the ViennaBiocenter, particularly Pawel Pasierbek, for support with microscopy; Mathias Madalinski for synthesizing the Ski7-P4 peptide; the VBCF NGS Unit (www.viennabiocenter.org/facilities) for RNA sequencing; Maria Novatchkova, Dominik Handler, and Brian Reichholf for helpful comments regarding RNA-seq data analyses; Karin Panser, Carina Pribitzer and the animal facility personnel for taking care of zebrafish; the VBC RNA Salon for helpful discussions and suggestions; the Pauli lab for valuable discussions and comments on the project and manuscript.

## Author contributions

L.E.C.Q. and A.P. designed the study. L.E.C.Q. performed the experiments with help from A.P. L.E.C.Q. analysed the experimental data. A.S. analysed the protein sequence conservation of Ski7. K.M. supervised mass-spectrometry analyses. L.E.C.Q. and A.P. wrote the manuscript with input from all authors.

## Data and materials availability

RNA-Seq data has been deposited at GEO (Gene Expression Omnibus) with accession code GSE147112.

## Code availability

Main scripts generated for the processing and analyses of data in this manuscript are available on GitHub (https://github.com/Quio-Enrique/pauliLab_Ski7_OET).

## Supplementary

### Sup. Data 1

Conservation analysis of Ski7 protein homologs.

### Sup. Data 2

Mass-spectrometric analysis of the Ski7-P4 peptide pull-down.

### Sup. Data 3

Mass-spectrometric analysis of TMT-MS of *WT* and *ski7^−/−^* embryos.

## References

Alexa A & Rahnenfuhrer J (2019) topGO: Enrichment Analysis for Gene Ontology. R package: 1–37

Anders S, Pyl PT & Huber W (2015) HTSeq--a Python framework to work with high-throughput sequencing data. Bioinformatics 31: 166–169

Araki Y, Takahashi S, Kobayashi T, Kajiho H, Hoshino S & Katada T (2001) Ski7p G protein interacts with the exosome and the Ski complex for 3‘-to-5’ mRNA decay in yeast. The EMBO Journal 20: 4684–4693

Bazzini AA, del Viso F, Moreno-Mateos MA, Johnstone TG, Vejnar CE, Qin Y, Yao J, Khokha MK & Giraldez AJ (2016) Codon identity regulates mRNA stability and translation efficiency during the maternal-to-zygotic transition. The EMBO Journal: e201694699–17

Benard L, Carroll K, Valle RC, Masison DC & Wickner RB (1999) The ski7 antiviral protein is an EF1-alpha homolog that blocks expression of non-Poly(A) mRNA in Saccharomyces cerevisiae. Journal of Virology 73: 2893–2900

Berner D, Zenkel M, Pasutto F, Hoja U, Liravi P, Gusek-Schneider GC, Kruse FE, Schödel J, Reis A & Schlötzer-Schrehardt U (2017) Posttranscriptional Regulation of LOXL1 Expression Via Alternative Splicing and Nonsense-Mediated mRNA Decay as an Adaptive Stress Response. Invest. Ophthalmol. Vis. Sci. 58: 5930–5940

Bertoldo MJ, Listijono DR, Ho W-HJ, Riepsamen AH, Goss DM, Richani D, Jin XL, Mahbub S, Campbell JM, Habibalahi A, Loh W-GN, Youngson NA, Maniam J, Wong ASA, Selesniemi K, Bustamante S, Li C, Zhao Y, Marinova MB, Kim L-J, et al (2020) NAD+ Repletion Rescues Female Fertility during Reproductive Aging. CellReports 30: 1670–1680.e8

Bray NL, Pimentel H, Melsted P & Pachter L (2016) Near-optimal probabilistic RNA-seq quantification. Nature Biotechnology: 1–4

Brunkard JO & Baker B (2018) A Two-Headed Monster to Avert Disaster: HBS1/SKI7 Is Alternatively Spliced to Build Eukaryotic RNA Surveillance Complexes. Front. Plant Sci. 9: 600–17

Bushnell B (2014) BBMap: A Fast, Accurate, Splice-Aware Aligner. Lawrence Berkeley National Laboratory: 1–3 Available at: https://escholarship.org/uc/item/1h3515gn

Carbone A, Zinovyev A & Kepes F (2003) Codon adaptation index as a measure of dominating codon bias. Bioinformatics 19: 2005–2015

Chew G-L, Pauli A, Rinn JL, Regev A, Schier AF & Valen E (2013) Ribosome profiling reveals resemblance between long non-coding RNAs and 5’ leaders of coding RNAs. Development 140: 2828–2834

Doblmann J, Dusberger F, Imre R, Hudecz O, Stanek F, Mechtler K & Dürnberger G (2019) apQuant: Accurate Label-Free Quantification by Quality Filtering. J. Proteome Res. 18: 535–541

Dorfer V, Pichler P, Stranzl T, Stadlmann J, Taus T, Winkler S & Mechtler K (2014) MS Amanda, a universal identification algorithm optimized for high accuracy tandem mass spectra. J. Proteome Res. 13: 3679–3684

Fujino Y, Yamada K, Sugaya C, Ooka Y, Ovara H, Ban H, Akama K, Otosaka S, Kinoshita H, Yamasu K, Mishima Y & Kawamura A (2018) Deadenylation by the CCR4- NOTcomplex contributes to the turnover of hairy-related mRNAs in the zebrafish segmentation clock. FEBS Letters 592: 3388–3398

Gagnon JA, Valen E, Thyme SB, Huang P, Ahkmetova L, Pauli A, Montague TG, Zimmerman S, Richter C & Schier AF (2014) Efficient Mutagenesis by Cas9 Protein-Mediated Oligonucleotide Insertion and Large-Scale Assessment of Single-Guide RNAs. PLoS ONE 9: e98186–8

Gauron C, Meda F, Dupont E, Albadri S, Quenech’Du N, Ipendey E, Volovitch M, Del Bene F, Joliot A, Rampon C & Vriz S (2016) Hydrogen peroxide (H2O2) controls axon pathfinding during zebrafish development. Developmental Biology 414: 133–141

Halbach F, Reichelt P, Rode M & Conti E (2013) The Yeast Ski Complex: Crystal Structure and RNA Channeling to the Exosome Complex. Cell 154: 814–826

Han Y, Ishibashi S, Iglesias-Gonzalez J, Chen Y, Love NR & Amaya E (2018) Ca2+-Induced Mitochondrial ROS Regulate the Early Embryonic Cell Cycle. CellReports 22: 218–231

He F, Celik A, Wu C & Jacobson A (2018) General decapping activators target different subsets of inefficiently translated mRNAs. eLife 7: 1–30

Hendrickson PG, Doráis JA, Grow EJ, Whiddon JL, Lim J-W, Wike CL, Weaver BD, Pflueger C, Emery BR, Wilcox AL, Nix DA, Peterson CM, Tapscott SJ, Carrell DT & Cairns BR (2017) Conserved roles of mouse DUX and human DUX4 in activating cleavage-stage genes and MERVL/HERVL retrotransposons. Nat Genetics 49: 925–934

Herberg S, Gert KR, Schleiffer A & Pauli A (2018) The Ly6/uPAR protein Bouncer is necessary and sufficient for species-specific fertilization. Science 361: 1029–1033

Horikawa W, Endo K, Wada M & Ito K (2016) Mutations in the G-domain of Ski7 cause specific dysfunction in non-stop decay. Sci Rep 6: 29295–10

Horvat F, Fulka H, Jankele R, Malik R, Jun M, Solcova K, Sedlacek R, Vlahovicek K, Schultz RM & Svoboda P (2018) Role of Cnot6lin maternal mRNA turnover. Life Sci. Alliance 1: e201800084–10

Houseley J & Tollervey D (2009) The Many Pathways of RNA Degradation. Cell 136: 763–776

Jamar NH, Kritsiligkou P & Grant CM (2017) The non-stop decay mRNA surveillance pathway is required for oxidative stress tolerance. Nucleic Acids Res 45: 6881–6893

Johnson AW & Kolodner RD (1995) Synthetic lethality of sep1 (xrn1) ski2 and sep1 (xrn1) ski3 mutants of Saccharomyces cerevisiae is independent of killer virus and suggests a general role for these genes in translation control. Mol. Cell. Biol. 15: 2719–2727

Kalisiak K, Kuliński TM, Tomecki R, Cysewski D, Pietras Z, Chlebowski A, Kowalska K & Dziembowski A (2016) A short splicing isoform of HBS1L links the cytoplasmic exosome and SKI complexes in humans. Nucleic Acids Res 23: gkw862–13

Katoh K, Misawa K, Kuma K-I & Miyata T (2002) MAFFT: a novel method for rapid multiple sequence alignment based on fast Fourier transform. Nucleic Acids Res 30: 3059–3066

Kim D, Paggi JM, Park C, Bennett C & Salzberg SL (2019) Graph-based genome alignment and genotyping with HISAT2 and HISAT-genotype. Nature Biotechnology: 1–12

Knoefler D, Thamsen M, Koniczek M, Niemuth NJ, Diederich A-K & Jakob U (2012) Quantitative In Vivo Redox Sensors Uncover Oxidative Stress as an Early Event in Life. Molecular Cell 47: 767–776

Kowalinski E, Kögel A, Ebert J, Reichelt P, Stegmann E, Habermann B & Conti E (2016) Structure of a Cytoplasmic 11-Subunit RNA Exosome Complex. Molecular Cell 63: 125–134

Kowalinski E, Schuller A, Green R & Conti E (2015) Saccharomyces cerevisiae Ski7 Is a GTP-Binding Protein Adopting the Characteristic Conformation of Active Translational GTPases. Structure/Folding and Design 23: 1336–1343

Labun K, Montague TG, Krause M, Torres Cleuren YN, Tjeldnes H & Valen E (2019) CHOPCHOP v3: expanding the CRISPR web toolbox beyond genome editing. Nucleic Acids Res 47: W171–W174

Lebreton A & Séraphin B (2008) Exosome-mediated quality control: substrate recruitment and molecular activity. Biochim. Biophys. Acta 1779: 558–565

Li H, Handsaker B, Wysoker A, Fennell T, Ruan J, Homer N, Marth G, Abecasis G, Durbin R1000 Genome Project Data Processing Subgroup (2009) The Sequence Alignment/Map format and SAMtools. Bioinformatics 25: 2078–2079

Link V, Shevchenko A & Heisenberg C-P (2006) BMC Developmental Biology. BMC Developmental Biology 6: 1–9

Liu J-J, Niu C-Y, Wu Y, Tan D, Wang Y, Ye M-D, Liu Y, Zhao W, Zhou K, Liu Q-S, Dai J, Yang X, Dong M-Q, Huang N & Wang H-W (2016) CryoEM structure of yeast cytoplasmic exosome complex. Cell Res 26: 822–837

Love MI, Huber W & Anders S (2014) Moderated estimation of fold change and dispersion for RNA-seq data with DESeq2. Genome Biol. 15: 31–21

Marshall AN, Han J, Kim M & van Hoof A (2018) Conservation of mRNA quality control factor Ski7 and its diversification through changes in alternative splicing and gene duplication. Proc. Natl. Acad. Sci. U.S.A. 115: E6808–E6816

Marshall AN, Montealegre MC, Jiménez-López C, Lorenz MC & van Hoof A (2013) Alternative Splicing and Subfunctionalization Generates Functional Diversity in Fungal Proteomes. PLoS Genet 9: e1003376–11

Mishima Y & Tomari Y (2016) Codon Usage and 3′ UTR Length Determine Maternal mRNA Stability in Zebrafish. Molecular Cell 61: 874–885

Mishima Y & Tomari Y (2017) Pervasive yet nonuniform contributions of Dcp2 and Cnot7 to maternal mRNA clearance in zebrafish. Genes Cells 22: 670–678

Mishima Y, Giraldez AJ, Takeda Y, Fujiwara T, Sakamoto H, Schier AF & Inoue K (2006) Differential Regulation of Germline mRNAs in Soma and Germ Cells by Zebrafish miR-430. Current Biology 16: 2135–2142

Montague TG, Cruz JM, Gagnon JA, Church GM & Valen E (2014) CHOPCHOP: a CRISPR/Cas9 and TALEN web tool for genome editing. Nucleic Acids Res 42: W401–W407

Nair S, Lindeman RE & Pelegri F (2012) In vitro oocyte culture-based manipulation of zebrafish maternal genes. Dev. Dyn. 242: 44–52

Nishida KM, Okada TN, Kawamura T, Mituyama T, Kawamura Y, Inagaki S, Huang H, Chen D, Kodama T, Siomi H & Siomi MC (2009) Functional involvement of Tudor and dPRMT5 in the piRNA processing pathway in Drosophila germlines. The EMBO Journal 28: 3820–3831

Orban TI & Izaurralde E (2005) Decay of mRNAs targeted by RISC requires XRN1, the Ski complex, and the exosome. RNA 11: 459–469

Pauli A, Norris ML, Valen E, Chew G-L, Gagnon JA, Zimmerman S, Mitchell A, Ma J, Dubrulle J, Reyon D, Tsai SQ, Joung JK, Saghatelian A & Schier AF (2014) Toddler: An Embryonic Signal That Promotes Cell Movement via Apelin Receptors. Science 343: 1248636–1248636

Pauli A, Valen E, Lin MF, Garber M, Vastenhouw NL, Levin JZ, Fan L, Sandelin A, Rinn JL, Regev A & Schier AF (2012) Systematic identification of long noncoding RNAs expressed during zebrafish embryogenesis. Genome Res. 22: 577–591

Petrova B, Liu K, Tian C, Kitaoka M, Freinkman E, Yang J & Orr-Weaver TL (2018) Dynamic redox balance directs the oocyte-to-embryo transition via developmentally controlled reactive cysteine changes. Proc. Natl. Acad. Sci. U.S.A. 115: E7978–E7986

Puigbò P, Bravo IG & Garcia-Vallve S (2008) CAIcal: A combined set of tools to assess codon usage adaptation. Biology Direct 3: 38–8

Quinlan AR & Hall IM (2010) BEDTools: a flexible suite of utilities for comparing genomic features. Bioinformatics 26: 841–842

Ritchie ME, Phipson B, Wu D, Hu Y, Law CW, Shi W & Smyth GK (2015) limma powers differential expression analyses for RNA-sequencing and microarray studies. Nucleic Acids Res 43: e47–e47

Schier AF (2007) The Maternal-Zygotic Transition: Death and Birth of RNAs. Science 316: 406–407

Schwanhäusser B, Busse D, Li N, Dittmar G, Schuchhardt J, Wolf J, Chen W & Selbach M (2011) Global quantification of mammalian gene expression control. Nature 473: 337–342

Selman K, Wallace RA, Sarka A & Qi X (1993) Stages of oocyte development in the zebrafish, Brachydanio rerio. J. Morphol. 218: 203–224

Smyth GK (2005) limma: Linear Models for Microarray Data. In Bioinformatics and Computational Biology Solutions Using R and Bioconductor, Gentleman R Carey VJ Huber W Irizarry RA & Dudoit S (eds) pp 397–420. New York, NY: Springer New York

Sohrabi-Jahromi S, Hofmann KB, Boltendahl A, Roth C, Gressel S, Baejen C, Soeding J & Cramer P (2019) Transcriptome maps of general eukaryotic RNA degradation factors. eLife 8: 1–29

Sorenson RS, Deshotel MJ, Johnson K, Adler FR & Sieburth LE (2018) ArabidopsismRNA decay landscape arises from specialized RNA decay substrates, decapping-mediated feedback, and redundancy. Proc Natl Acad Sci USA 115: E1485–E1494

Subtelny AO, Eichhorn SW, Chen GR, Sive H & Bartel DP (2014) Poly(A)-tail profiling reveals an embryonic switch in translational control. Nature 508: 66–71

Sun M, Schwalb B, Pirkl N, Maier KC, Schenk A, Failmezger H, Tresch A & Cramer P (2013) Global Analysis of Eukaryotic mRNA Degradation Reveals Xrn1-Dependent Buffering of Transcript Levels. Molecular Cell 52: 52–62

Tadros W & Lipshitz HD (2009) The maternal-to-zygotic transition: a play in two acts. Development 136: 3033–3042

Taus T, Köcher T, Pichler P, Paschke C, Schmidt A, Henrich C & Mechtler K (2011) Universal and confident phosphorylation site localization using phosphoRS. J. Proteome Res. 10: 5354–5362

Toh-E A, Guerry P & Wickner RB (1978) Chromosomal superkiller mutants of Saccharomyces cerevisiae. J. Bacteriol. 136: 1002–1007

Tomecki R, Kristiansen MS, Lykke-Andersen SOR, Chlebowski A, Larsen KM, Szczesny RJ, Drazkowska K, Pastula A, Andersen JS, Stepien PP, Dziembowski A & Jensen TH (2010) The human core exosome interacts with differentially localized processive RNases: hDIS3 and hDIS3L. The EMBO Journal 29: 2342–2357

van Hoof A, Frischmeyer PA, Dietz HC & Parker R (2002) Exosome-mediated recognition and degradation of mRNAs lacking a termination codon. Science 295: 2262–2264

van Hoof A, Staples RR, Baker RE & Parker R (2000) Function of the ski4p (Csl4p) and Ski7p proteins in 3‘-to-5’ degradation of mRNA. Mol. Cell. Biol. 20: 8230–8243

Vandaele L, Thys M, Bijttebier J, Van Langendonckt A, Donnay I, Maes D, Meyer E & Van Soom A (2010) Short-term exposure to hydrogen peroxide during oocyte maturation improves bovine embryo development. Reproduction 139: 505–511

Wang L, Lewis MS & Johnson AW (2005) Domain interactions within the Ski2/3/8 complex and between the Ski complex and Ski7p. RNA 11: 1291–1302

Waterhouse AM, Procter JB, Martin DMA, Clamp M & Barton GJ (2009) Jalview Version 2--a multiple sequence alignment editor and analysis workbench. Bioinformatics 25: 1189–1191

Weick E-M, Puno MR, Januszyk K, Zinder JC, DiMattia MA & Lima CD (2018) Helicase-Dependent RNA Decay Illuminated by a Cryo-EM Structure of a Human Nuclear RNA Exosome-MTR4 Complex. Cell 173: 1663–1677.e21

Wolf J & Passmore LA (2014) mRNA deadenylation by Pan2–Pan3. Biochemical Society Transactions 42: 184–187

Wu D & Dean J (2020) EXOSC10 sculpts the transcriptome during the growth-to-maturation transition in mouse oocytes. bioRxiv: 663377

Yu C, Ji S-Y, Sha Q-Q, Dang Y, Zhou J-J, Zhang Y-L, Liu Y, Wang Z-W, Hu B, Sun Q-Y, Sun S-C, Tang F & Fan H-Y (2016) BTG4 is a meiotic cell cycle–coupled maternal-zygotic-transition licensing factor in oocytes. Nat Struct Mol Biol 23: 387–394

