## Supplementary_Figures for "Zebrafish Ski7 tunes RNA levels during the oocyte-to-embryo transition"

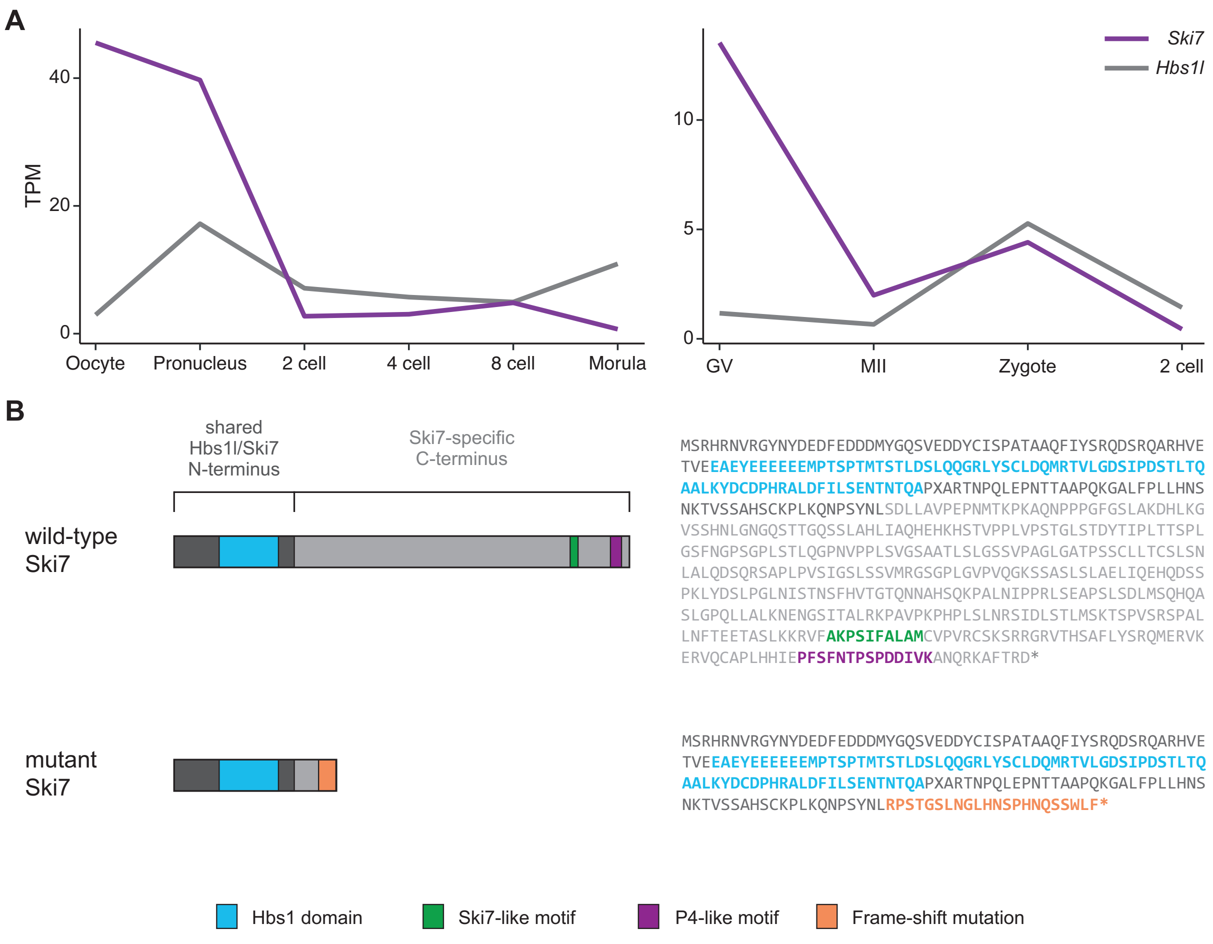


**Sup. Fig. 1. Murine *Ski7* RNA expression and protein sequence of wild-type and mutant zebrafish Ski7.**
A) Expression levels (TPM) of mouse *Ski7* (purple) and *Hbs1l* (grey) during early embryogenesis. Left: *Hendrickson PG, et al 2017*; right: *Yu C, et al, 2016*.
B) Zebrafish mutant Ski7 protein lacks the conserved Ski7-like and P4-like motifs. Schematic (left) and amino acid sequence (right) of wild-type and mutant Ski7 proteins. Specific domains and motifs are highlighted. The mutant protein contains the shared N-terminus from Hbs1l but lacks the Ski7-like and P4-like motifs due to a premature stop codon.

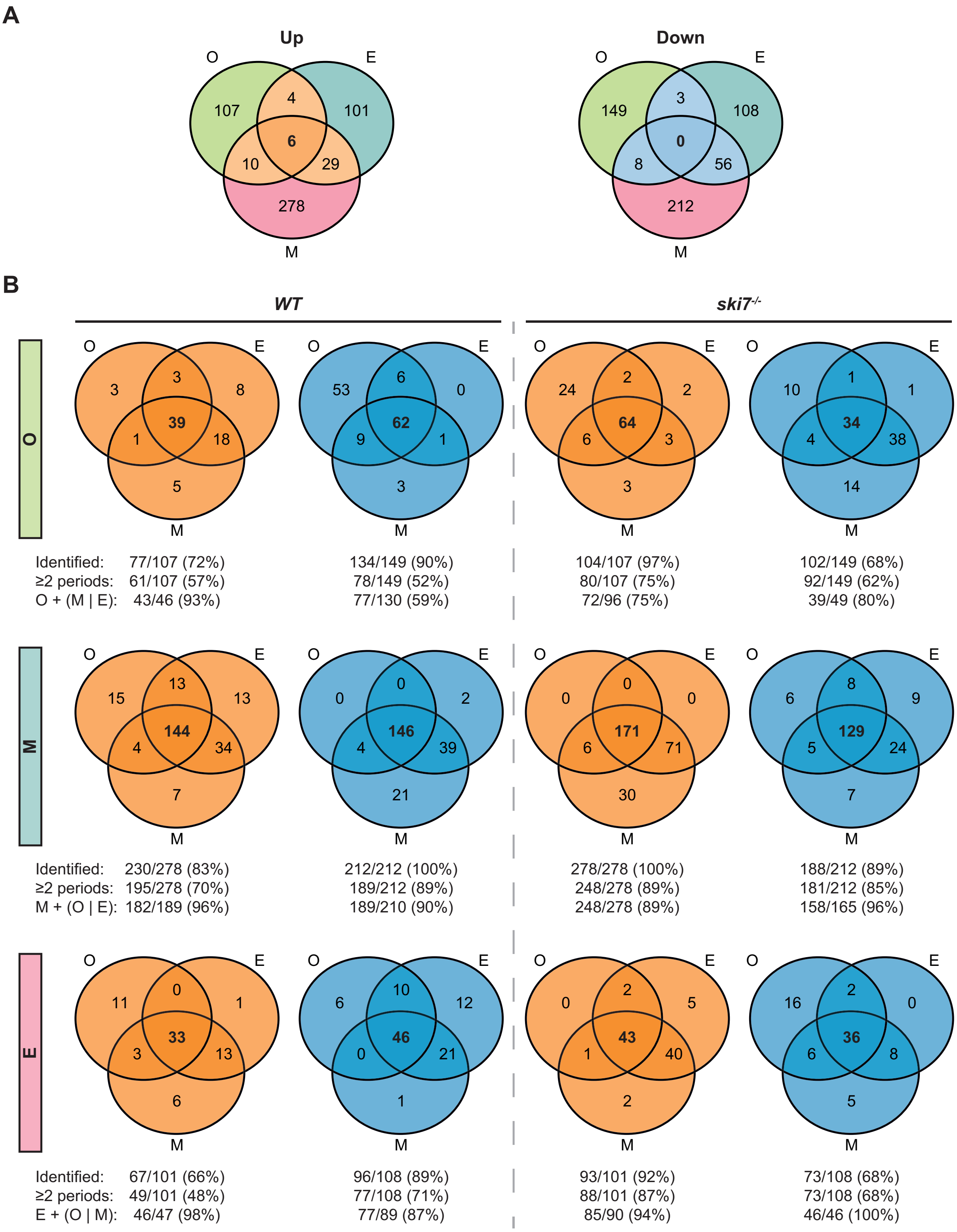
A
Up
O
E
107
4
101
6
10
29
278
M
Down
O
E
149
3
108
0
8
56
212
M
B
WT
ski7-/-
O
O
E
3
3
8
39
1
18
5
M
O
E
53
6
0
62
9
1
3
M
O
E
24
2
2
64
6
3
3
M
O
E
10
1
1
34
4
38
14
M
Identified: 77/107 (72%)
≥2 periods: 61/107 (57%)
O + (M | E): 43/46 (93%)
134/149 (90%)
78/149 (52%)
77/130 (59%)
104/107 (97%)
80/107 (75%)
72/96 (75%)
102/149 (68%)
92/149 (62%)
39/49 (80%)
M
O
E
15
13
13
144
4
34
7
M
O
E
0
0
2
146
4
39
21
M
O
E
0
0
0
171
6
71
30
M
O
E
6
8
9
129
5
24
7
M
Identified: 230/278 (83%)
≥2 periods: 195/278 (70%)
M + (O | E): 182/189 (96%)
212/212 (100%)
189/212 (89%)
189/210 (90%)
278/278 (100%)
248/278 (89%)
248/278 (89%)
188/212 (89%)
181/212 (85%)
158/165 (96%)
E
O
E
11
0
1
33
3
13
6
M
O
E
6
10
12
46
0
21
1
M
O
E
0
2
5
43
1
40
2
M
O
E
16
2
0
36
6
8
5
M
Identified: 67/101 (66%)
≥2 periods: 49/101 (48%)
E + (O | M): 46/47 (98%)
96/108 (89%)
77/108 (71%)
77/89 (87%)
93/101 (92%)
88/101 (87%)
85/90 (94%)
73/108 (68%)
73/108 (68%)
46/46 (100%)

**Sup. Fig. 2. Targets of Ski7 are broadly expressed but regulated at a specific time during the oocyte-to-embryo transition.**
A) Venn diagrams of the overlap of DEGs from the three periods from Fig. 3F.
B) Venn diagrams of the overlap of 'period-specific' DEGs (identified in all stages of a given period as differentially regulated) during the oocyte-to-embryo transition in *WT* and *ski7*$^{-/-}$ samples. The numbers below refer to the expression in wild type (left) or *ski7*$^{-/-}$ (right). Identified = expressed in wild type or *ski7*$^{-/-}$ in at least one period; ≥2 periods = expressed in wild type or *ski7*$^{-/-}$ in at least two periods; X + Y|Z = expressed in the period of origin (X) and at least one other period (Y and/or Z).

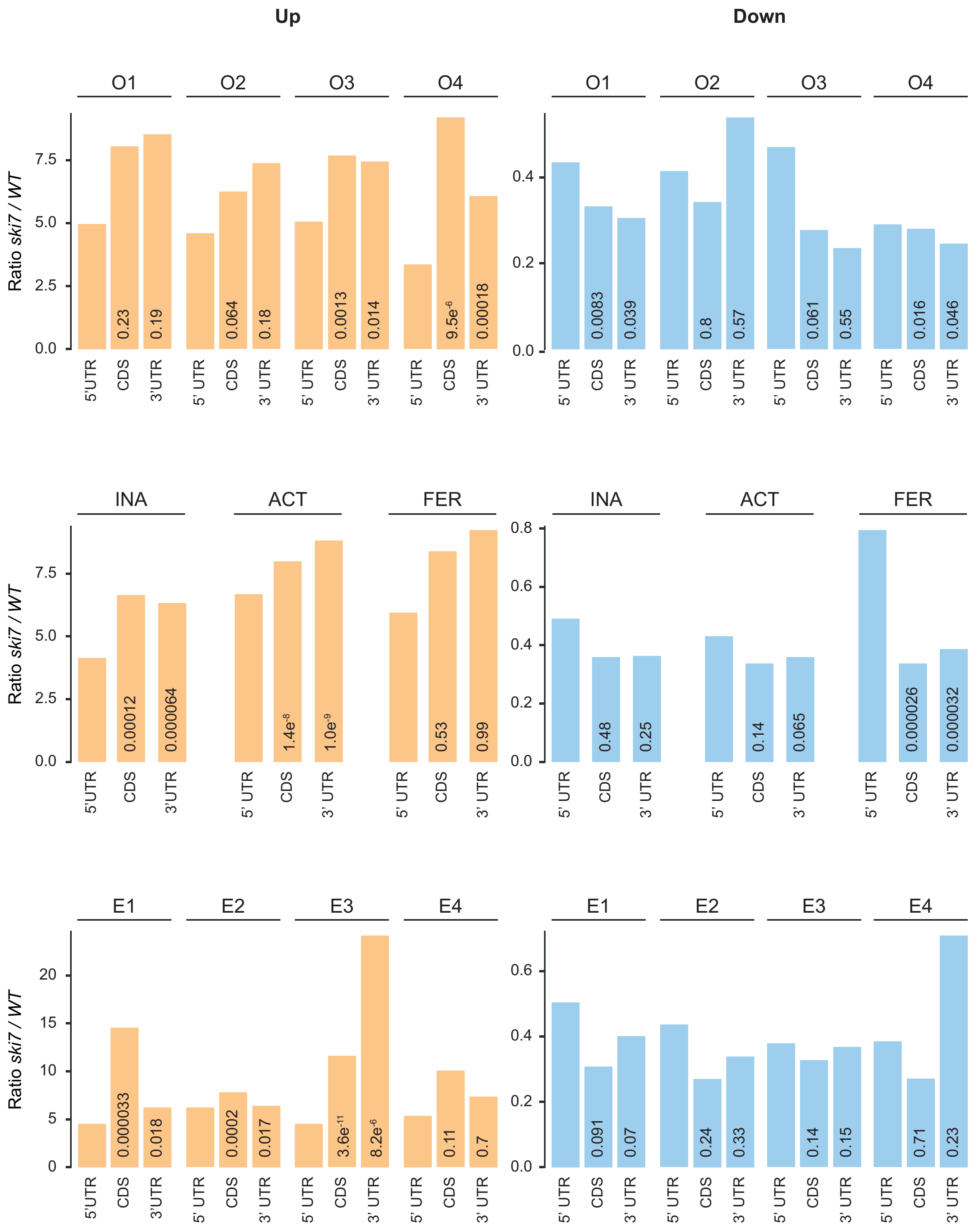


**Sup. Fig. 3. Genes upregulated in *ski7*$^{-/-}$ mutants show higher read accumulation towards their 3' ends.**

Ratio of RNA-seq read densities (*ski7*$^{-/-}$ / *wild-type*) across different transcript regions 5' UTR, CDS, 3' UTR). Up-regulated genes: orange; down-regulated genes: blue. The numbers in the bars show the p-values (Wilcoxon test) for the comparison of the corresponding CDS or 3' UTR against the 5'UTR.

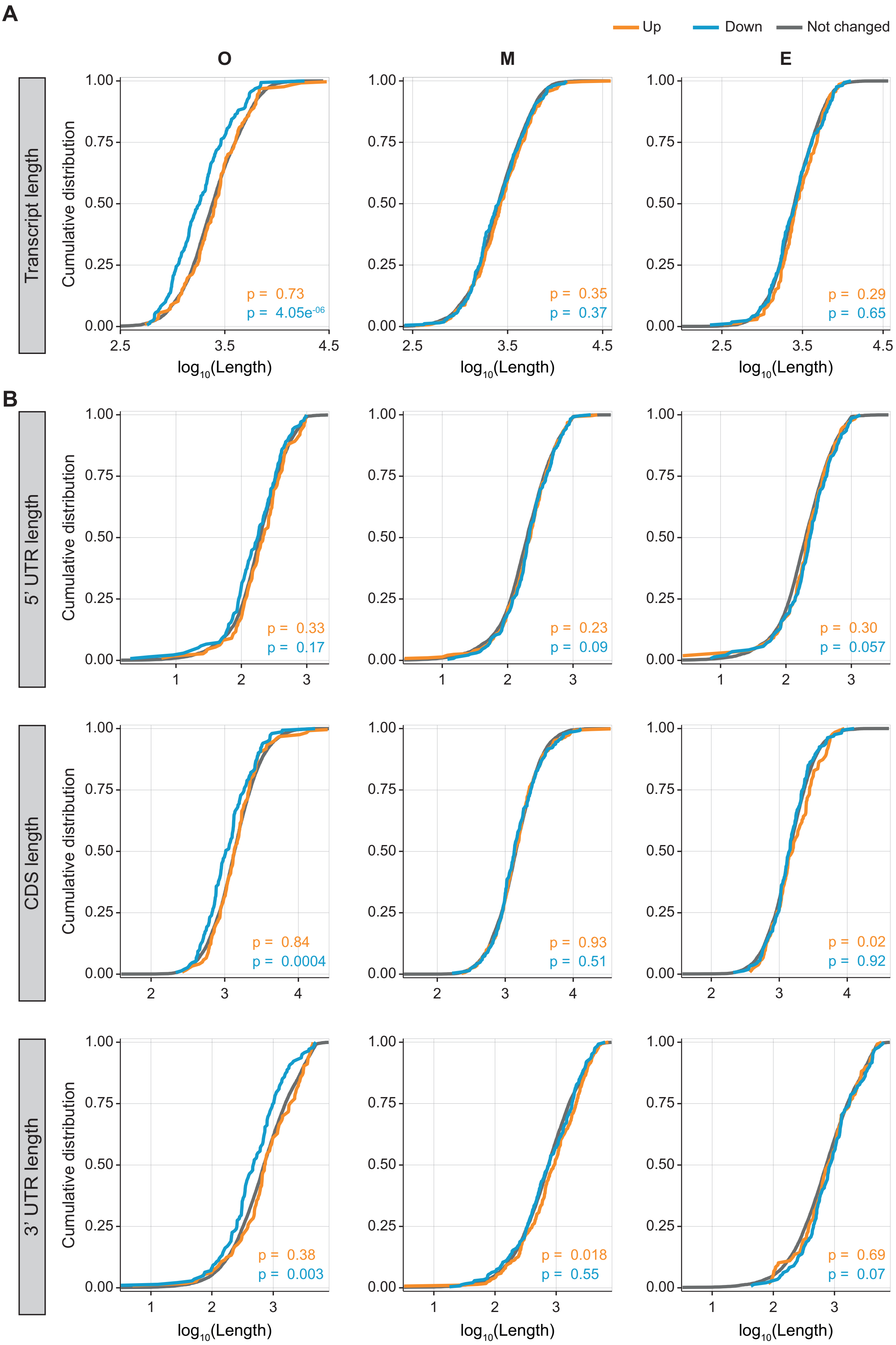

A
B
Up
Down
Not changed
O
M
E
Transcript length
5' UTR length
CDS length
3' UTR length
Cumulative distribution
log10(Length)
p = 0.73
p = 4.05e-06
p = 0.35
p = 0.37
p = 0.29
p = 0.65
p = 0.33
p = 0.17
p = 0.23
p = 0.09
p = 0.30
p = 0.057
p = 0.84
p = 0.0004
p = 0.93
p = 0.51
p = 0.02
p = 0.92
p = 0.38
p = 0.003
p = 0.018
p = 0.55
p = 0.69
p = 0.07

**Sup. Fig. 4. Oogenesis genes down-regulated in the absence of Ski7 tend to be shorter.**
Cumulative distribution of transcript lengths of up-regulated (orange), down-regulated (blue) and unchanged (grey) genes per period.
A) Analyses of the length of full transcripts (Kolmogorov-Smirnov p = $4.05e^{-6}$).
B) Analyses of the length per transcript region (5' UTRs, CDSes and 3' UTRs). Note that the biggest difference in down-regulated genes during oogenesis is observed in CDSes and 3' UTRs (Kolmogorov-Smirnov p = 0.0004 & 0.003, respectively).

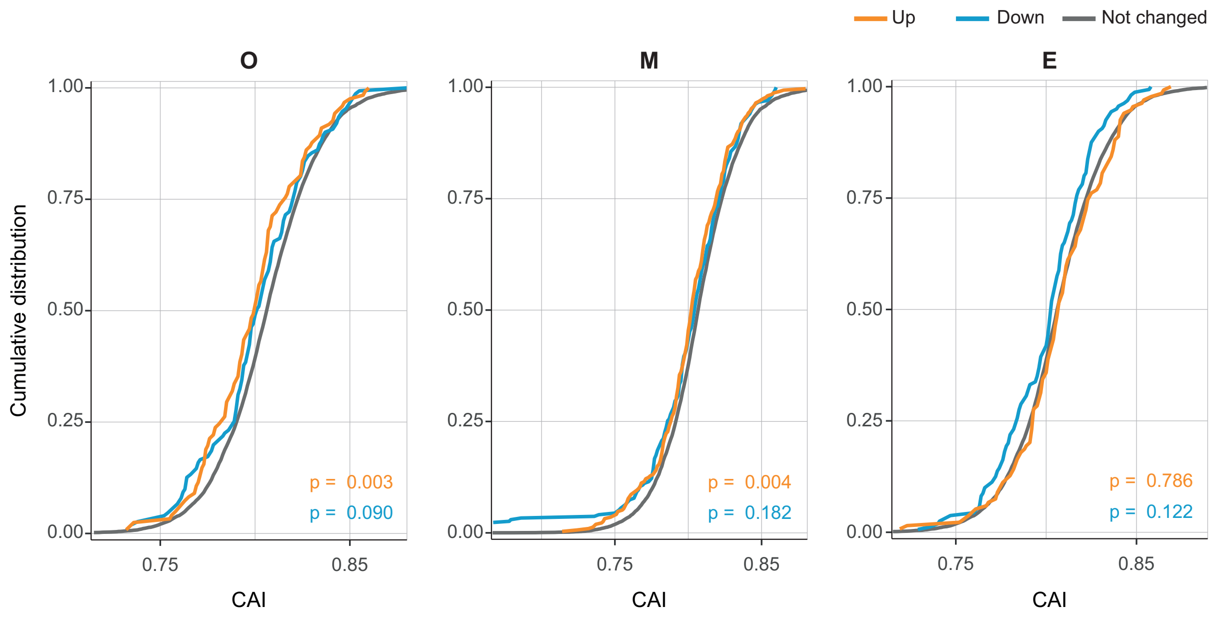


**Sup. Fig. 5. Genes up-regulated in *ski7*$^{-/-}$ mutants during oogenesis and in the egg are enriched for rare codons.**

Cumulative fraction of the usage of synonymous codons in DEGs (up-regulated: orange, down-regulated: blue) and unchanged genes (grey) per period as measured by codon adaptation index (CAI). P-values from Kolmogorov-Smirnov test.

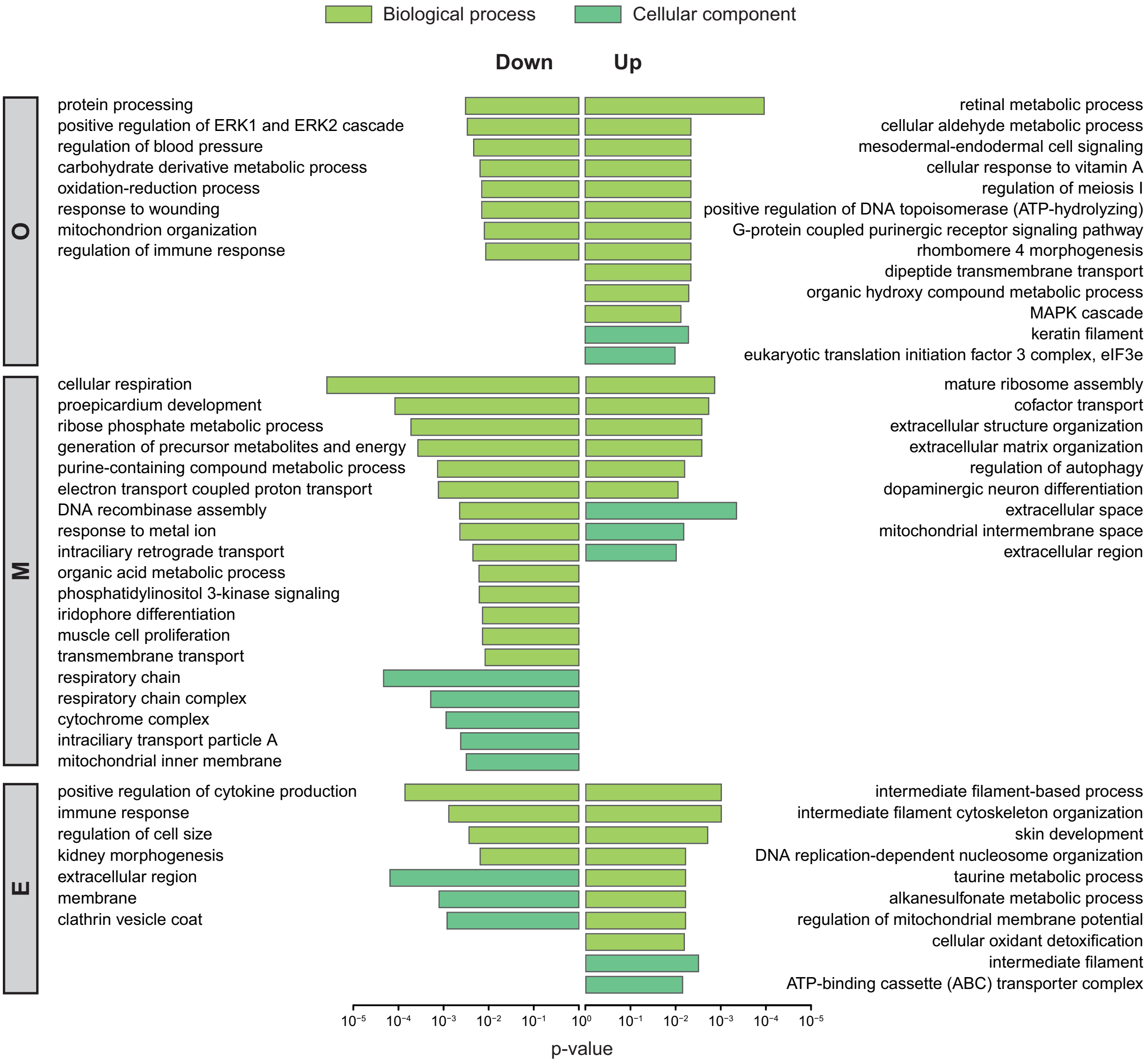


**Sup. Fig. 6. GO terms enriched in differentially expressed genes in *ski7*$^{-/-}$ mutants.**

GO terms enriched in the overlapping set of DEGs from each period are shown for the categories 'biological process' (light green) and 'cell compartment' (dark green) (related to Fig 5A).

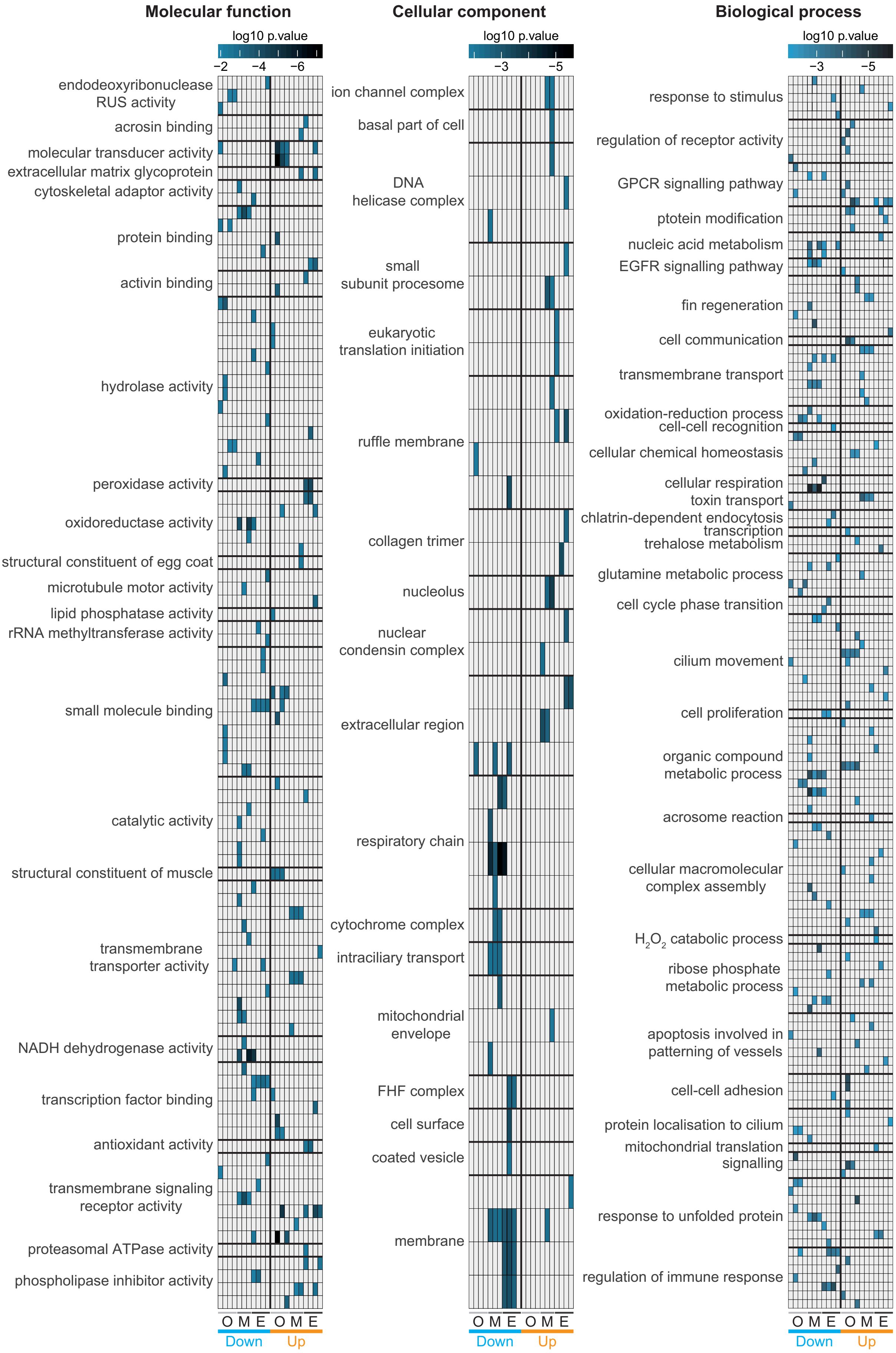

Molecular function
log10 p.value
−2 −4 −6
endodeoxyribonuclease RUS activity
acrosin binding
molecular transducer activity
extracellular matrix glycoprotein
cytoskeletal adaptor activity
protein binding
activin binding
hydrolase activity
peroxidase activity
oxidoreductase activity
structural constituent of egg coat
microtubule motor activity
lipid phosphatase activity
rRNA methyltransferase activity
small molecule binding
catalytic activity
structural constituent of muscle
transmembrane transporter activity
NADH dehydrogenase activity
transcription factor binding
antioxidant activity
transmembrane signaling receptor activity
proteasomal ATPase activity
phospholipase inhibitor activity
O M E O M E
Down Up
Cellular component
log10 p.value
−3 −5
ion channel complex
basal part of cell
DNA helicase complex
small subunit procesome
eukaryotic translation initiation
ruffle membrane
collagen trimer
nucleolus
nuclear condensin complex
extracellular region
respiratory chain
cytochrome complex
intraciliary transport
mitochondrial envelope
FHF complex
cell surface
coated vesicle
membrane
O M E O M E
Down Up
Biological process
log10 p.value
−3 −5
response to stimulus
regulation of receptor activity
GPCR signalling pathway
ptotein modification
nucleic acid metabolism
EGFR signalling pathway
fin regeneration
cell communication
transmembrane transport
oxidation-reduction process
cell-cell recognition
cellular chemical homeostasis
cellular respiration
toxin transport
chlatrin-dependent endocytosis
transcription
trehalose metabolism
glutamine metabolic process
cell cycle phase transition
cilium movement
cell proliferation
organic compound metabolic process
acrosome reaction
cellular macromolecular complex assembly
H2O2 catabolic process
ribose phosphate metabolic process
apoptosis involved in patterning of vessels
cell-cell adhesion
protein localisation to cilium
mitochondrial translation
signalling
response to unfolded protein
regulation of immune response
O M E O M E
Down Up

**Sup. Fig. 7. GO terms enriched in differentially expressed genes for each stage across the oocyte-to-embryo transition.**

GO terms enriched in DEGs at each stage of the time-course experiment during oocyte-to-embryo transition. O = oogenesis, M = mature eggs, E = embryogenesis.

**A**

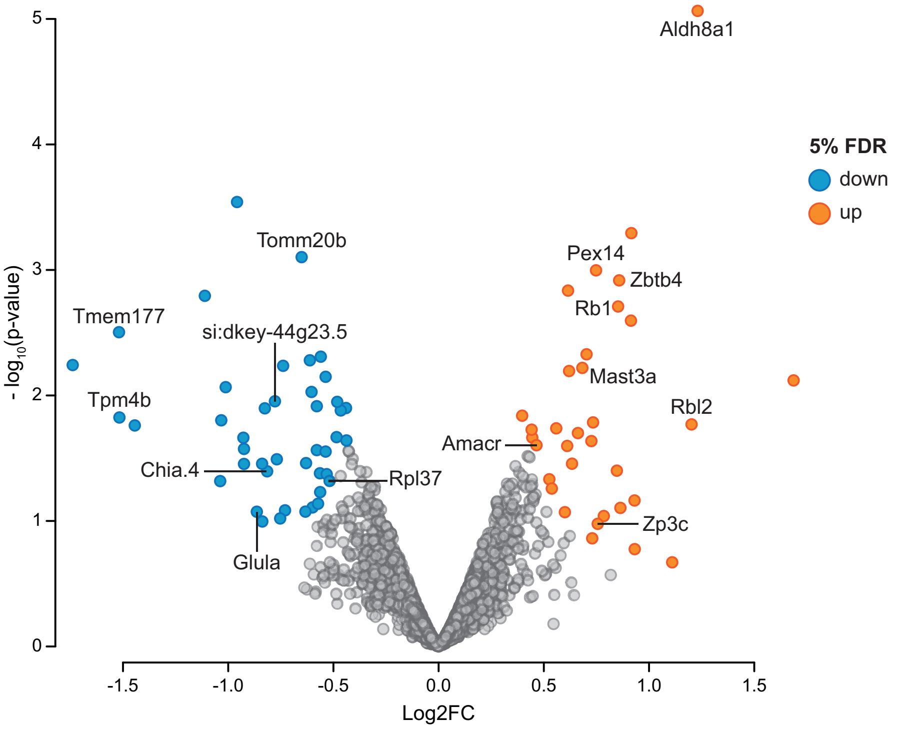


**B**

| Gene name | Annotated function |
|---|---|
| *fga* | fibrinogen complex |
| *fgb* | fibrinogen complex |
| *fgg* | fibrinogen complex |
| *ccar2* | transcription and splicing |
| *nrip1b* | transcription regulation |
| *myg1* | nuclear localization |
| *nr2f6b* | transcriptional repression |
| ***tomm20b*** | **import into mathochondrial matrix** |
| *ninl* | calcium ion binding activity |
| *itln3* | ——— |
| *ccdc9* | ——— |
| *dhx34* | 3'-5' RNA helicase activity |
| ***glula*** | **glutamate-ammonia ligase activity** |
| ***rpl37*** | **rRNA binding activity** |
| ***tpm4b*** | **actin filament binding activity** |
| *ube2j1* | ubiquitin conjugating enzyme activity |
| *ociad2* | complement system |
| *plg* | blood coagulation and tissue remodeling |
| *atp5mea* | ATP synthesis |
| *c6* | complement system |
| *mgat5* | synthesis of glycoprotein oligosaccharides |
| *cfhl5* | complement factor H like 5 |
| *ahsg1* | negative regulator of endopeptidase activity |
| *fetub* | negative regulator of endopeptidase activity |
| *rab5if* | integral component of membrane |
| ***chia.4*** | **chitin catabolic process** |
| *rca2.1* | complement system |
| *nptna* | cell-cell interaction |
| *lmod2b* | tropomyosin binding activity |
| *cd9b* | lateral line development |
| *ndufb2* | respiratory chain complex I |
| *aldh2.1* | aldehyde dehydrogenase activity |
| *padi2* | vasculature development |
| *fbxo22* | ubiquitin-protein transferase activity |
| ***tmem177*** | **cytochrome c oxidase maturation** |
| *ppp2r3b* | protein dephosphorylation |
| *pdk2a* | pyruvate dehydrogenase kinase |
| ***si:dkey-44g23.5*** | **orthologous to human MCRIP2** |
| *si:ch211-145c1.1* | ——— |
| *si:ch211-210c8.6* | oxidoreductase activity |
| *si:ch211-126j24.1* | ion channel binding activity |
| *si:ch211-212c13.10* | endopeptidase inhibitor activity |
| *si:ch211-207i20.3* | DNA binding activity |

| Gene name | Annotated function |
|---|---|
| *micall2b* | metal ion binding activity |
| *lipt1* | protein lipoylation |
| ***zp3c*** | **egg coat formation** |
| *ckmt2a* | creatine kinase, mitochondrial 2a |
| *vps26bl* | intracellular protein transport |
| ***zbtb4*** | **transcription factor activity** |
| *lhpp* | protein dephosphorylation |
| *gsta.2* | glutathione transferase activity |
| *dph1* | biosynthesis of diphthamide |
| *slc18b1* | transmembrane transport |
| *znf395a* | nucleic acid binding activity |
| ***mast3a*** | **cytoskeleton organization** |
| *rbp1* | retinoic acid biosynthetic process |
| ***amacr*** | **racemase** |
| ***rb1*** | **activating TF binding activity** |
| *kdelc1* | protein O-linked glycosylation |
| *kdelc2* | protein O-linked glycosylation |
| *fitm2* | fat storage |
| *pfkma* | phosphofructokinase |
| ***pex14*** | **protein import into peroxisome matrix** |
| ***aldh8a1*** | **NAD-dependent oxidation** |
| *sept15* | cytoskeletal GTPase |
| ***rbl2*** | **activating TF binding activity** |
| *llph* | RNA Pol II binding activity |
| *ugt2a4* | UDP-glucuronosyltransferase |
| *rtf2* | replication termination at RTS1 barrier |
| *dock11* | guanyl-nucleotide exchange factor activity |
| *usp12b* | thiol-dependent ubiquitinyl hydrolase activity |
| *cep78* | centrosome localization |
| *chdh* | choline dehydrogenase activity |
| *noc2l* | negative regulation of transcription |
| *si:dkey-97o5.1* | ——— |
| *si:ch211-133n4.7* | carbohydrate binding activity |

**Sup. Fig. 8. Differentially expressed proteins in *ski7*$^{-/-}$ embryos.**
A) Volcano plot of proteins identified by tandem mass tag mass spectrometry (TMT-MS) from wild-type and *ski7*$^{-/-}$ mutant embryos at 4 hours post-fertilization. Significantly up- and down-regulated proteins are coloured in orange and blue, respectively. Labelled dots highlight proteins for which the mRNA was also identified as differentially regulated by RNA-seq.
B) List of all significantly up- and down-regulate proteins (indicated by gene name) from the TMT-MS analyses. Differentially expressed proteins for which the mRNA was also identified as differentially expressed by RNA-seq are highlighted in bold. Genes that have been associated with stress response or oxidative/reductive stress are highlighted in green.
